# Species identification and genotyping of *Citrobacter* spp. using genes with high nucleotide diversity

**DOI:** 10.1101/2025.10.28.685243

**Authors:** Nobuyoshi Yagi, Rin Nashiro, Itaru Hirai

## Abstract

*Citrobacter* spp. are facultative anaerobic Gram-negative bacillus found in a wide range of habitats. Recently, there have been reports of an increasing number of cases of *Citrobacter* spp. being the cause of nosocomial infections, and of increasing highly antimicrobial-resistant (AMR) *Citrobacter* spp. In clinical laboratory testing, *Citrobacter* spp., such as *C. freundii* and *C. braakii*, are handled collectively as the *C. freundii* complex. This can be obstacle to collect information regarding species identification and genetic lineage which are necessary for estimating distribution mechanism and evaluating AMR degree for each *Citrobacter* species. This study investigated gene combinations that allow for *Citrobacter* spp. identification and genetic lineage estimation with as few genes as possible. *Citrobacter* genomes of 18 species were collected as many as possible from GenBank. After the gene annotation, nucleotide diversity of 1,113 genes contained in all 453 genomes was calculated. Genotypes of top seven highest nucleotide diversity (HND) genes were confirmed and investigated their distributions among *Citrobacter* genomes. Except for *C. werkmanii* and *C. cronae*, it was suggested that the genotype of the top HND gene, groups_3152, and combinations of the genotypes of the top two HND genes were a species- and genetic lineage-specific distribution, respectively. Considering that the genomes of *C. werkmanii* and *C. cronae* were highly similar and indistinguishable each other by conventional genotyping methods, it was suggested that the combination of the top two HND genes could be used to *Citrobacter* spp. identification and genetic lineage classification.

**Importance Statement:** Molecular epidemiology of individual *Citrobacter* species has not been widely performed, partly because *Citrobacter* spp. are handled as the *C. freundii* complex in the clinical testing and the multi-locus sequencing typing (MLST) has been performed targeting the entire genus *Citrobacter*. This study proposed a combination of two genes for identification of *Citrobacter* spp. and classify their genetic lineages. Currently, phylogenetic tree analysis has become possible using core-genome MLST and core-genome single nucleotide polymorphisms (SNPs) after whole genome sequencing (WGS) of bacterial isolates. But these phylogenetic analyses require a considerable amount of cost. In this regard, using two genes for *Citrobacter* species identification and genetic lineage classification reduces the required cost compared to WGS, encouraging its introduction into clinical testing and further performing molecular epidemiology of individual *Citrobacter* species.

## Introduction

Members of genus *Citrobacter* are facultative anaerobic Gram-negative bacillus that are detected in human, animal and environmental samples. Some species of *Citrobacter*, including *C. freundii*, are known to cause nosocomial and opportunistic infections such as urinary tract infections, meningitidis and sepsis [1-4].

Regarding nosocomial infections, *Citrobacter* spp. have been regarded as one of the important causing agents [5-7]. A systematic review written by Fonton, Hassoun-Kheir and Harbarth indicated that number of hospitalized-patients with *Citrobacter* infection an outbreak frequency of *Citrobacter* outbreak in among hospitalized-patients were increasing [8]. Antibiotics treatment of infections caused by *C. freundii*, that is a main causative species, has sometimes been unsuccessful because *C. freundii* carries inducible AmpC β-beta lactamase gene, and induced AmpC β-lactamase nullifies β-lactams including the third generation cephalosporins [9-11]. Currently higher AMR bacteria, such as extended-spectrum β-lactamase (ESBL)-producing and carbapenem-resistant *Citrobacter* spp., have emerged and spread in many countries [1, 12-14]. It is not difficult to understand that the emerging of these AMR bacteria makes treatments of infections caused by *Citrobacter* spp. increasingly difficult. It reminds that *Citrobacter* spp. should be tracked regularly and sufficiently to see if it causes infections or develops further multidrug resistance by surveillance or monitoring. In addition, it is important to identify the transmission route and evaluate clonality of these causing clinical isolates in nosocomial infections from the perspective of infection control. However, *Citrobacter* spp. such as *C. braakii, C. freundii, C. gillenii, C. murliniae, C. rodentium, C. sedlakii, C. werkmanii* and *C. youngae* are often handled as the *C. freundii* complex, and the species and clones of bacterial strain causing nosocomial infection have not always been identified, which hinders the estimation of the transmission route of the causative strains and the evaluation of their clonality.

Genetic characteristics, such as phylogeny, antimicrobial-resistance genes (ARGs), pathogenic genes, plasmid replicons, etc., have been collected from bacterial strains obtained from nosocomial infections or outbreaks. Because, these genetic characteristics are essential information for surveillance and have been subjected to estimate the clonality and potential transmission routes of these pathogenic bacterial strains. Several methods, such as pulsed-field gene electrophoresis (PFGE) [15], multi-locus sequence typing (MLST) [16], MALDI-TOF mass spectrometry (MS) [17] and WGS [18, 19] have been used for collecting the genetic characteristics of pathogenic and/or antimicrobial-resistant clinical isolates.

Regarding *Citrobacter* spp., it was reported that there are certain atypical strains in the most clinically important *C. freundii* [20]. The presence of atypical *C. freundii* strains, that could make it difficult to identify species using standard microbiological biochemical tests, might affect the selection of candidate test isolates for PFGE. Regarding the MLST, in the conventional MLST scheme used for *Citrobacter* spp., it is known that some *Citrobacter* strains may have gene deletion or major mutations in genes that are used in the MLST scheme, which could sometimes cause problems such as the inability to determine their sequence types (STs) [21, 22].

Because of these analytical methods’ disadvantages, currently WGS and the following bioinformatics with obtaining bacterial genome information has been the most widely used method. However, performing WGS needs a certain amount of expense and labor, it has not become a routine examination procedure for pathogenic bacterial surveillance in healthcare-associated facilities yet. Currently, confirming the clonality of pathogenic bacterial isolates and estimation of the transmission routes of causing clinical isolates have been performed mainly as research studies [23-25].

It is highly possible that WGS of pathogenic and AMR bacteria will be introduced into clinical testing in the future. Therefore, in this study, assuming the introduction of WGS into clinical testing, we investigated whether there would be a gene combination that could identify *Citrobacter* spp. of pathogenic and AMR bacteria with a minimum number of genes and make up for the shortcomings of conventional MLST. And for this purpose, we considered the application of nucleotide diversity (π) to gene selection. Specifically, we collected 453 *Citrobacter* genomes from the public database GenBank and extracted core genes commonly detected in all collected genomes. The extracted core genes were ranked by nucleotide diversity and the top seven highest nucleotide diversity (HND) genes were examined. Our results suggested that the top two HND genes were sufficient to identify *Citrobacter* spp. Furthermore, it was suggested that the genetic relatedness of *Citrobacter* strains was assessable by displaying them on a phylogenetic tree drawn based on core genome SNPs by using a notation system based on genotypes of the top two HND genes and the ST of each genome.

## Materials and Methods

### *Citrobacter* genomes and species confirmation

Genomes of genus *Citrobacter* were retrieved from the GenBank database [26] (finally accessed on Aug 24th, 2024) as shown in **Supplementary Table S1**. Reference genomes for each *Citrobacter* spp. that are registered in RefSeq [27] were designated as the reference dataset for FastANI analysis [28]. Each genome of *Citrobacter* spp. was compared against the reference dataset. *Citrobacter* spp. name of the reference genome showed the highest average nucleotide identity (ANI) and more than 95 % was assigned as *Citrobacter* spp. of each genome sequence. MLST was performed on each *Citrobacter* genome to determine their STs at the PubMLST website [29].

### Core genes extraction

Gene annotation of all collected *Citrobacter* genomes was performed using Prokka [30], and the resulting annotations were used to construct a pangenome database with PanTA [31]. Genes, *i*.*e*., ORF, potentially producing hypothetical proteins that were identified by Prokka were grouped by PanTA and indicated as “groups_1”, “groups_2”, “groups_3” and so forth in the order in which they were detected. Among the genes in the pangenome database, genes detected in all *Citrobacter* genomes, target core genes, core genes, soft core genes, shell genes and cloud genes were defined as genes detected in 100%, 99.0% or more but less than 100%, 95.0% or more but less than 99.0%, 15.0% or more but less than 95.0%, 0% or more but less than 15.0% of the *Citrobacter* genomes, respectively.

### Calculating nucleotide diversity of core genome genes and genotyping using top seven HND genes

Nucleotide diversity of the target core genes was calculated using R and RStudio with pegas and ape packages. The target core genes were sorted in descending order of nucleotide diversity, and the top seven HND genes were subjected to a clustering analysis by using the VSEARCH software [32]. In this study, it was set that sequences classified into the same genotype should have 100% sequence similarity between sequences. These consequence sequences of the top seven HND genes were sorted in descending order of detection frequency and numbered. Genotypes of the top seven HND genes contained in each genome were confirmed using the BLAST [33].

### Phylogenetic tree drawing

The top seven HND genes of the *Citrobacter* genomes were aligned using MAFFT [34]. Phylogenetic inference was then carried out using VeryFastTree [35, 36], and the resulting tree was visualized in iTOL [37]. In addition, the top seven HND genes were aligned and the top HND gene and concatenated sequences of the HND genes were used for phylogenetic reconstruction. The phylogenetic tree visualization was also performed with VeryFastTree and iTOL.

The core gene was identified by using Prokka and PanTA as mentioned above. Then, approximately-maximum-likelihood phylogenetic tree was constructed by using VeryFastTree with GTR+CAT model and 1,000 bootstrap replicates. Finally, resulting tree was visualized by using iTOL.

## Results

In this study, 453 *Citrobacter* genomes were collected from the GenBank and bacterial species of the 453 genomes belonging to 18 *Citrobacter* spp. were identified (**Supplementary Table S1**). Among them, 178 (39.3%) genomes were *C. freundii* and followed by *C. braakii* (81 genomes, 17.9%) and *C. portucalensis* (55 genomes, 12.1%) as shown in **Table 1**. Regarding ST, STs were determined species-specifically in 13 (72.2%) of the 18 *Citrobacter* spp. No identical ST was not observed in two or more than two *Citrobacter* spp. In total, 132 types of STs were determined in 304 (67.1%) of 453 genomes. The detected STs of *Citrobacter* spp. were shown in **Table 1** and **Supplementary Table S1** and **S2**.

**Table 1.** Genomes examined in this study.

| Table 1. Genomes examined in this study. |  |  |  |  |  |  |  |
| --- | --- | --- | --- | --- | --- | --- | --- |
| Species | N | % | MLST |  |  |  | Major STs |
|  |  |  | Determined |  | N.D. |  |  |
|  |  |  | N | % | N | % |  |
| *C. freundii* | 178 | 39.3 | 157 | 88.2 | 21 | 11.8 | 22(31), 116(12), 150(8) |
| *C. braakii* | 81 | 17.9 | 48 | 59.3 | 33 | 40.7 | 603(9), 110(7), 1250(5) |
| *C. portucalensis* | 55 | 12.1 | 35 | 63.6 | 20 | 36.4 | 143(4), 252(3), 17(2) |
| *C. koseri* | 35 | 7.7 | 25 | 71.4 | 10 | 28.6 | 690(8), 862(4), 854(3) |
| *C. gillenii* | 30 | 6.6 | 4 | 13.3 | 26 | 86.7 | 1115(2), 699(1), 1251(1) |
| *C. amalonaticus* | 18 | 4.0 | 12 | 66.7 | 6 | 33.3 | 710(3), 712(3), 716(3) |
| *C. cronae* | 18 | 4.0 | 8 | 44.4 | 10 | 55.6 | 31(3), 259(3), 108(1) |
| *C. werkmanii* | 8 | 1.8 | 4 | 50.0 | 4 | 50.0 | 104(3), 108(1) |
| *C. youngae* | 8 | 1.8 | 4 | 50.0 | 4 | 50.0 | 128(1), 218(1), 420(1), 969(1) |
| *C. farmeri* | 3 | 0.7 | 3 | 100.0 | 0 | 0.0 | 1007(2), 686(1) |
| *C. sedlakii* | 3 | 0.7 | 2 | 66.7 | 1 | 33.3 | 942(2) |
| *C. europaeus* | 3 | 0.7 | 1 | 33.3 | 2 | 66.7 | 1341(1) |
| *C. meridianamericanus* | 3 | 0.7 | 1 | 33.3 | 2 | 66.7 | 1066(1) |
| *C. pasteurii* | 3 | 0.7 | 0 | 0.0 | 3 | 100.0 | - |
| *C. rodentium* | 3 | 0.7 | 0 | 0.0 | 3 | 100.0 | - |
| *C. enshiensis* | 2 | 0.4 | 0 | 0.0 | 2 | 100.0 | - |
| *C. arsenatis* | 1 | 0.2 | 0 | 0.0 | 1 | 100.0 | - |
| *C. tructae* | 1 | 0.2 | 0 | 0.0 | 1 | 100.0 | - |
| Total | 453 | 100 | 304 | 67.1 | 149 | 32.9 |  |

Genes contained in each *Citrobacter* genome were confirmed (**Supplementary Table S1 and S3**). In the 453 *Citrobacter* genomes analyzed, in total 57,628 genes were detected and 4635.2 ± 292.2 (mean ± S.D.) genes were contained in one genome on average. The median of the detected gene number was 4644, and the maximum and the minimum detected gene number was 7757 and 3731, respectively. Among the detected genes, 1,113 (1.9%) genes, 1,357 (2.4%) genes, 316 (0.5%) genes, 2,307 (4.0%) genes and 52,535 (91.2%) genes were classified as target core genes, core genes, soft core genes, shell genes and cloud genes, respectively (**Supplementary Table S3**).

Then, nucleotide diversity of the 1,113 target core genes was calculated (**Supplementary Table S4**). The nucleotide diversity of the target core genes ranged from 0.0041 to 0.1808. The average number of the nucleotide diversity of the target core genes was 0.0785 ± 0.0284 (mean ± S.D.), and the median was 0.0797. The calculated nucleotide diversity of the target core genes was indicated by the box plot. There were seven genes above the upper whisker, such as groups_3152 (0.1808), *nanK* (0.1782), *iprA* (0.1723), *mipA* (mipA_2, 0.1644), *yehY* (0.1556), *yhcH* (yhcH_2, 0.1552) and *ymdB* (0.1521). In this study, we evaluated whether these top seven HND genes were applicable for species identification and classification of *Citrobacter* spp. as shown in **Fig. 1**.

**Fig. 1.**
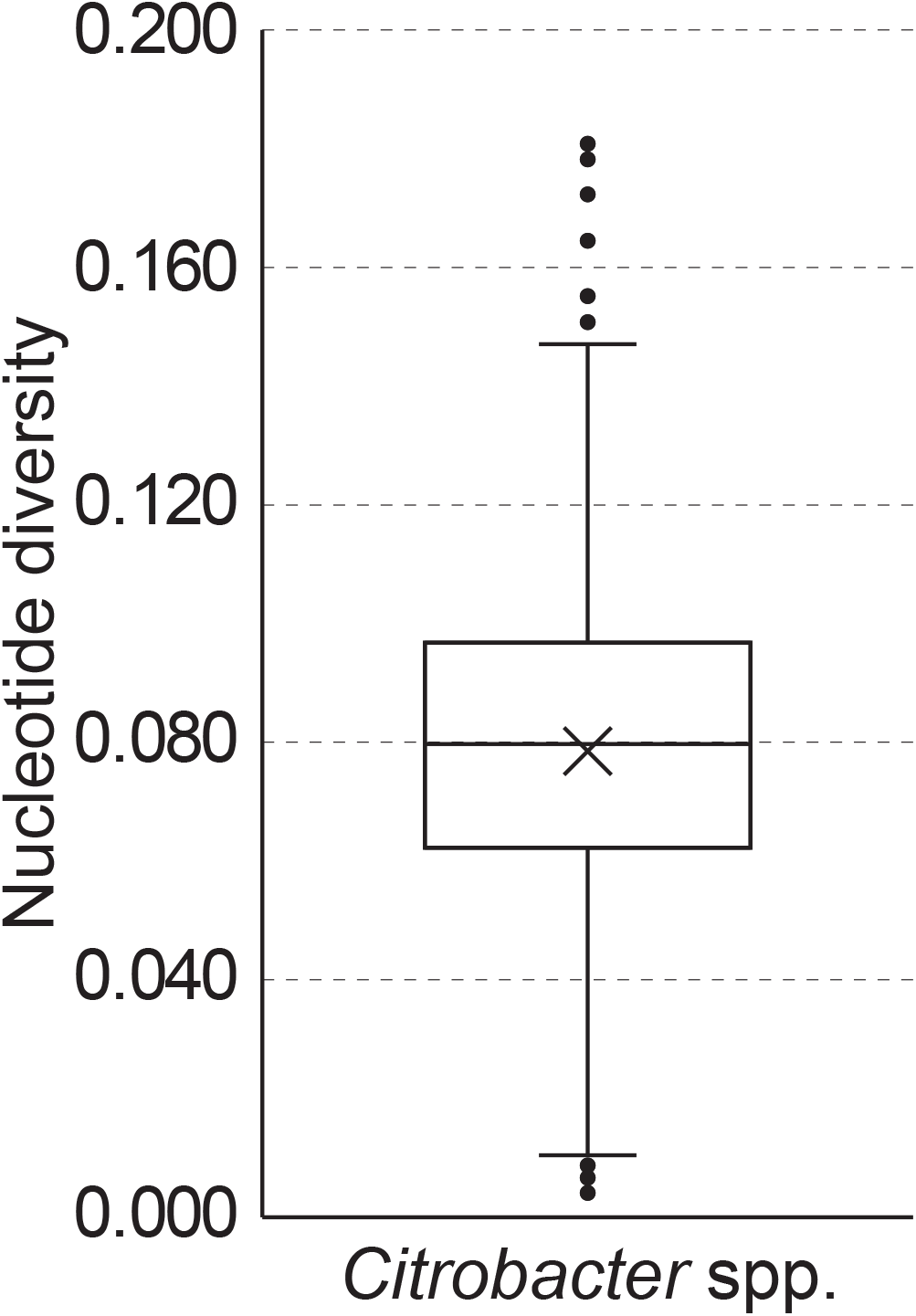
A box plot of nucleotide diversity. Nucleotide diversity of the 1,113 target core genes was calculated. There are seven genes indicating higher nucleotide diversity above the upper whisker. These seven HND genes were, groups_3152, *nanK, iprA, mipA* (mipA_2), *yehY, yhcH* (yhcH_2) and *ymdB* and their nucleotide diversity rates were 0.1808, 0.1782, 0.1723, 0.1644, 0.1556, 0.1552 and 0.1521, respectively.

Genotypes of the top seven HND genes were observed, and 161, 216, 170, 153, 218, 149 and 164 kinds of genotypes were confirmed in groups_3152, *nanK, iprA, mipA, yehY, yhcH* and *ymdB*, respectively. Based on this, genotypes of the top seven HND genes in each of the 453 *Citrobacter* genomes were confirmed (**Supplementary Table S1**). There were 237, 255, 262, 271, 272 and 275 combinations of genotypes of the top two, three, four, five, six and seven HND genes. Then, it was confirmed whether the genotypes of the top seven HDN genes were distributed among *Citrobacter* spp. in a species-specific manner. It seemed that even only the genotypes of the top HND gene, groups_3152, appeared to be distributed species-specifically among *Citrobacter* spp. Even though, there were the exceptions of three genotypes, such as groups_3152-13, -43 and -56, commonly observed in *C. cronae* and *C. werkmanii* (**Supplementary Table S5**). Combinations of the top two HND genes, *i*.*e*., groups_3152 and *nanK*, were distributed species-specifically among *Citrobacter* spp. except for one exception, *i*.*e*., groups_3152-43 and *nanK*-70 (**Supplementary Table S6**). There were two genomes containing the groups_3152-43 and nanK-70 genotypes, and one was *C. cronae* (Genome # 317 in **Supplementary Table S1**) and the other was *C. werkmanii* (Genome # 413 in **Supplementary Table S1**). ST of these two genomes was not determined by the PubMLST and observed genotypes of the top seven HND genes were perfectly matched. Therefore, even using genotypes of the top seven HND genes could not discriminate between these two genomes.

To observe distribution of *Citrobacter* genomes on a phylogenetic tree, concatenated sequences of the top two HND genes were used to draw phylogenetic trees (HND tree). For comparison, a phylogenetic tree based on core genome SNPs (cgSNP tree) was drawn (**Fig. 2**). Genomes of major *Citrobacter* spp. such as *C. freundii, C. braakii, C. portucalensis, C. koseri, C. gillenii* and *C. amalonaticus* were distributed on separate branches on the cgSNP tree. However, the detected cgSNPs of *C. cronae* and *C. werkmanii* genomes were not enough to locate genomes of these two *Citrobacter* spp. on separate branches, consequently, these two *Citrobacter* spp. genome were located on the same branch (**Fig. 2A**). Interestingly, *Citrobacter* spp. were well separately distributed on separate branches of the HND tree as same manner as the cg SNP tree (**Fig. 2B**). However, genomes of the two *Citrobacter* spp., *C. cronae* and *C. werkmanii*, were again located on the same branch.

**Fig. 2.**
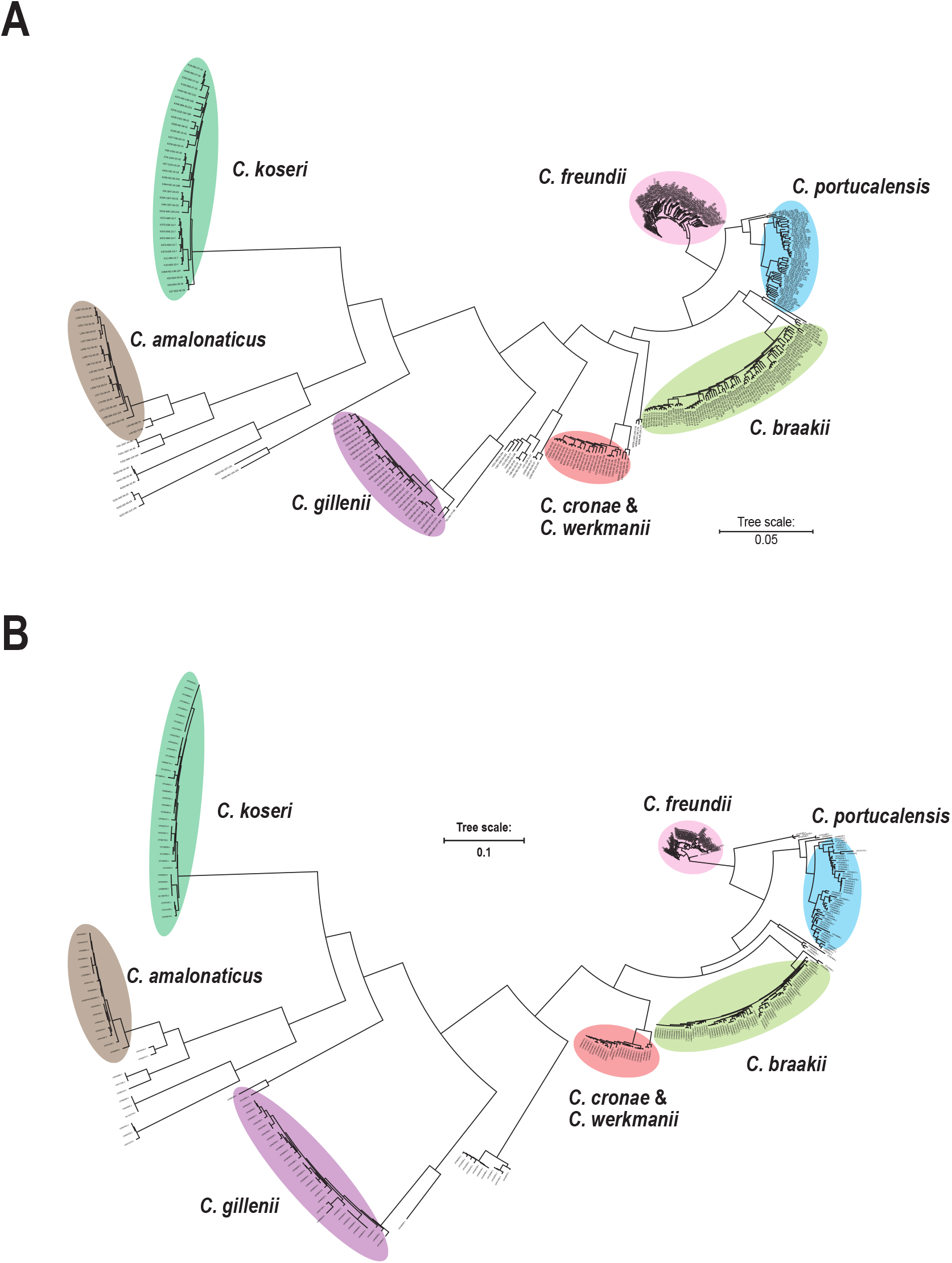
Distribution of *Citrobacter* spp. genomes on phylogenetic trees. (**A**) A phylogenetic tree based on core genome SNPs were drawn. (**B**) A phylogenetic tree based on concatenated sequences of the top two HND genes were drawn. Major *Citrobacter* spp. are shown with color indicators. *C. freundii*, light red, *C. braakii*, yellow green, *C. portucalensis*, blue, *C. koseri*, green, *C. gillenii*, pink, *C. amalonaticus*, brown, *C. cronae* and *C. werkmanii*, orange.

To understand the relationship between the genotypes of the top two HND genes and ST of each genome and the distribution on the cgSNP tree, close observation of the branches of the cgSNP tree was performed (**Supplementary Fig. S1-S17**). To do this, each genome was indicated by a coding rule consisted of one letter notation of *Citrobacter* spp., serial number of the genome examined, the determined ST, genotypes of the top two HND genes (**Supplementary Fig. S18**). Generally, genomes with the same ST and same genotypes of the top two HND genes were closely located in a branch (or rather a twig). In addition, there were three manners of genome arrangement of *Citrobacter* genomes on the cgSNP tree, such as, 1) genomes with undetermined ST but located near genome(s) with certain ST on the cgSNP tree, 2) genomes located near genome(s) on the cgSNP tree but their ST and/or genes in the top seven HND genes were different and 3) genomes with the same ST (or both were ND) and located near genome(s) on the cgSNP tree but their genes in the top 7 HND genes were different (**Table 2**).

**Table 2.** Summary of observation.

**1. Genomes with undetermined ST but located near genome(s) with certain ST on the cgSNP tree.**
| Coding # of genome | Coding # used for comparison | Numerical indication of MLST scheme genes | Difference of genes in MLST scheme | Numerical indication of the top 7 HND genes | Difference of the top 7 HND genes<sup>1)</sup> | Presumed ST |
| --- | --- | --- | --- | --- | --- | --- |
| D106-ND-1-1 | D17-22-1-1 | 5-15-12-11-20-0-15 | *lysP* deletion | 1-1-2-3-1-1-1 | none | 22 |
| D129-ND-9-13 | D140-270-9-13 | 5-10-12-0-7-11-6 | *dnaG* deletion | 9-13-8-49-10-1-23 | *mipA* | 270 |
| D133-ND-9-13 | D140-270-9-13 | 5-10-12-0-7-11-6 | *dnaG* deletion | 9-13-8-49-10-1-23 | *mipA* | 270 |
| D351-ND-14-10 | D446-18-14-10 | 0-10-14-5-17-14-12 | *arcA* deletion | 14-10-6-2-189-14-16 | *yehY* | 18 |
| K401-ND-15-18 | K67-1154-15-18 | 101-214-0-299-293-178-293 | *clpX* deletion | 15-18-26-24-20-30 | none | 1154 |
| C36-ND-20-81 | C208-31-20-22 | 11-0-18-20-27-15-21 | *aspC* deletion | 20-81-15-19-14-17 | *nanK* mutation (G813A) | - |
| C206-ND-20-22 | C208-31-20-22 | 11-24-0-20-27-15-21 | *clpX* deletion | 20-22-15-19-14-17 | none | 31 |
| L251-ND-25-67 | L427-563-25-67 | 115-137-0-184-178-185-194 | *clpX* deletion | 25-67-20-15-73-66-12 | none | 563 |
| D358-ND-12-38 | D221-65-12-38 | 5-16-17-11-20-74-0 | *mdh* deletion | 12-38-1-138-191-2-148 | *iprA* mutations (in D221, G403T and deleted after C405) | - |
| D444-ND-1-2 | D34-655-5-80 | 5-28-167-9-33-5-6 | *lysP* mutations (T406C, A441T, C444T) | 1-2-6-5-48-1-9 | Groups_3152 mutations (G13A, G135T, G201A, T220C, T276A, G354T, C429G) and *nanK* mutations (16 nucleotides) | - |

**2. Genomes located near genome(s) on the cgSNP tree but their ST and/or genes in the top seven HND genes were different.**
| Coding # of genome | Coding # used for comparison | Numerical indication of MLST scheme genes | Difference of genes in MLST scheme | Numerical indication of the top 7 HND genes | Difference of the top 7 HND genes<sup>1)</sup> |
| --- | --- | --- | --- | --- | --- |
| D53-1095-3-2 | D390-8-3-2 | 5-5-336-5-7-5-6 | *clpX* mutation (C342T) | 3-2-1-1-11-1-10 | none |
| D29-257-8-4 | D74-95-8-4 | 5-51-53-6-33-114-15 | *lysP* mutation (T552C) | 8-4-1-10-5-4-20 | *ymdB* |
| D128-167-9-13 | D140-270-9-13 | 5-10-12-11-7-11-72 | *mdh* mutation (A382G) | 9-13-8-1-10-1-23 | none |
| D153-519-11-4 | D156-481-11-4 | 7-127-10-6-13-11-13 | *mdh* mutation (G303A) | 11-4-56-1-65-5-63 | none |
| D340-112-36-72 | D28-622-36-79 | 5-26-5-54-69-17-15 | *fadD* mutation (G1176A) | 36-72-1-6-44-9-38 | *nanK* mutation (T672A) |
| D389-1098-28-1 | D345-118-28-1 | 5-64-333-22-33-14-6 | *clpX* mutation (G841T) | 28-1-37-1-3-1-29 | none |

| Coding # of genome | Coding # used for comparison | Numerical indication of MLST scheme genes | Difference of genes in MLST scheme | Numerical indication of the top 7 HND genes | Difference of the top 7 HND genes<sup>1)</sup> |
| --- | --- | --- | --- | --- | --- |
| D349-22-1-1 | D17-22-1-1 | 5-15-12-11-20-17-15 | none | 1-1-67-3-16-1-1 | *iprA* mutation (G423A) and *yehY* |
| D165-150-5-118 | D24-150-5-2 | 10-10-12-9-13-11-10 | none | 5-2-12-2-22-1-2 | *nanK* (C763T) and *iprA* (deletion and mutations after G384) |
| D259-95-8-4 | D197-95-8-4 | 5-51-53-6-33-5-15 | none | 8-4-1-114-5-4-8 | *mipA* |
| D41-98-11-12 | D44-98-11-12 | 7-54-55-6-60-11-52 | none | 11-12-78-1-9-9-18 | *iprA* mutation (A251G) |
| D42-98-11-12 | D44-98-11-12 | 7-54-55-6-60-11-52 | none | 11-12-1-73-9-9-18 | *mipA* |
| D101-169-21-99 | D79-169-21-19 | 7-80-10-6-56-61-13 | none | 21-99-27-1-15-1-11 | Deletion in *nanK* (from A250 to A270) |
| B173-959-31-34 | B148-959-31-115 | 30-138-140-125-58-147-49 | none | 31-34-10-32-26-27-46 | *nanK* mutation (T482C) |
| B81-ND-38-53 | B299-ND-38-53 | 20-148-0-97-102-208-81 | none | 38-53-28-43-53-50-40 | *nanK* mutation (G164T) |

<sup>1)</sup> Mutated nucleotide positions are indicated only in top 3 HND genes.

## Discussion

In this study, top seven HND genes were selected from the 1,113 target core genes extracted from the *Citrobacter* spp. genomes that were collected from GenBank as many as possible. Among the top seven HND genes, the top two HND genes, Groups_3152 and *nanK*, were evaluated whether these genes were applicable to identify *Citrobacter* spp. and to discriminate genetic lineages.

As WGS of bacterial strains has been widely performed, it is becoming easier to obtain whole genome information of bacterial strains. Even though, some species of *Citrobacter* genus are handled as *C. freundii* complex. Our results showed that the distribution of *Citrobacter* genomes on the HND phylogenetic tree drawn based on the concatenated sequences of the top two HND genes was similar manner to that observed in the cgSNP tree (**Fig. 2**), suggesting that the two HND genes could identify *Citrobacter* species and showed similar classification power for genetic lineages as cgSNP. In other words, our results suggested that the top two HND genes, *i*.*e*., genotypes of these two genes, were applicable to identify *Citrobacter* spp. except for *C. werkmanii* and *C cronae*, such as *C. freundii, C. braakii, C. portucalensis, C. koseri, C. gillenii*, and *C. amalonaticus* by considering the distribution of *Citrobacter* genomes on the phylogenetic trees (**Fig. 2**). Identification of the species of *Citrobacter freundii* complex would enable us to evaluate the progression of AMR in bacterial strains and to consider the distribution routes of nosocomial-infections for each bacterial species more precisely.

Regarding *C. werkmanii* and *C. cronae*, distributions of these two species’ genomes were not separate each other not only in HND tree but also in cgSNP tree (**Fig. 2**). To confirm this, we calculated the ANI using the genomes of these two bacterial species, and % ANI values were higher than 96%, suggesting that it was not easy to classify these two bacterial species by genome analysis (**Supplementary Table S7**). Consistent with our calculations, study, that reported *C. cronae* as a novel species, indicated %ANI between genomes of the novel *C. coronae* isolate and *C. werkmani*i NBRC105721 was 95.9–96% and only way to separate these two *Citrobacter* spp. were digital DNA–DNA hybridization [38]. Therefore, it was indicated that other methods for classifying these two bacterial species should be needed and that these two species might be handled for instance such as *C. werkmanii* complex.

We observed some *Citrobacter* genomes that were located on the same branch (or twig) in the cgSNP tree, but whose STs could not be determined or whose STs were different (**Supplementary Fig. S1-S18**). Our results indicated that these genomes either lacked one of the genes in the MLST scheme or were due to gene mutations (mainly single point mutations in a single gene) (**Table 2, Supplementary Fig. S1-S18**). Furthermore, if the genotypes of the top two HND genes in these genomes were the same, they were located on the same twig of the cgSNP tree, suggesting that they share the same genetic background. In other words, it was suggested that the genotypes of the top two HND genes could be used to presume undetermined STs and evaluate the genetic relatedness between two different STs.

As *Escherichia coli* ST131, ST is useful because it can be used like a proper noun when indicating genetic lineage. However, in the case of the genus *Citrobacter*, MLST is performed on the entire genus, and the ST number has already exceeded 1,000. It made it inconvenient to use it like a proper noun and might have reduced its practicality.

In this regard, it was suggested that the genotype combinations of the top two HND genes were applicable to classification of *Citrobacter* spp. precisely. Because the genotype combinations of the top two HND genes used in this study generally showed species-specific distributions, and the only exception was a combination of groups_3152-43 and nanK-70, which were detected in *C. werkmanii* and *C. cronae* (**Supplementary Table S6**). Therefore, as an instance, a coding rule that combines the one-letter notation of *Citrobacter* spp. and the genotypes of the top two HND genes with or without ST indicating *Citrobacter* spp. and genetic lineage could be useful for molecular epidemiology and estimation of transmission routes.

In the future, WGS of clinical isolates may be introduced in clinical laboratories [39, 40]. Because data analysis including phylogenetic analysis using the obtained genetic information in WGS requires a certain amount of effort, it might be a huge burden for clinical testing. Even to construct a cgSNP tree, there are several steps, such as obtaining sequence data from WGS, obtaining whole genome sequences by *de novo* assembly or reference mapping, selecting core genes from the analyzing and already published data, extracting SNPs in the core genes and constructing phylogenetic trees. In the genotyping of the top two HND genes, obtaining whole genome sequence can be directly applied to BLAST-based analysis against the genotype databases to identify *Citrobacter* spp. and genotypes of the top two HND genes that could be useful to presume genetic lineage of the analyzed isolates. In this respect, it was suggested that this study could contribute to reducing the effort required for data analysis after WGS even in the clinical laboratories.

As described above, this study described a new method for *Citrobacter* spp. identification and phylogenetic analysis that could complement the weaknesses of MLST. However, there are at least two limitations in this study. First, while this study used all collectable *Citrobacter* genomes that could be obtained as whole genomes from GenBank, the database used in this study is species-biased, and the number of genomes analyzed for minor species is insufficient. Our preliminary consideration suggested that the sequences of the top two HND genes used in this study vary significantly among the *Citrobacter* spp. However, it cannot be ruled out that the ranking of HND genes may change if genomes of these minor *Citrobacter* spp. are further included. Second, when WGS are performed and the resulting established whole genomes are annotated, it has been confirmed that the length of the sequence recognized as the coding sequence (CDS) differs. This may also affect the ranking of HND genes. Therefore, to address this issue, it is recommended that CDSs should be determined by reference mapping for the annotation after WGS. In this study, we only performed analysis using genome data, we could not confirm this issue by actual WGS of *Citrobacter* spp. isolates.

## Data Availability Statement

Data available on request from the authors

## Acknowledgements

This work was partly supported by the KAKENHI 23K09672 to IT and 24K20197 to NY.

