## Supplementary Figures for "Species identification and genotyping of *Citrobacter* spp. using genes with high nucleotide diversity"

**Supplementary Figure S1-18**

Supplementary Figure S1: Coding rule for *Citrobacter* spp. genomes.
Supplementary Figure S2: Distribution of genomes containing a genotype, groups_3152-1.
Supplementary Figure S3: Distribution of genomes containing a genotype, groups_3152-2.
Supplementary Figure S4: Distribution of genomes containing a genotype, groups_3152-3.
Supplementary Figure S5: Distribution of genomes containing a genotype, groups_3152-4.
Supplementary Figure S6: Distribution of genomes containing a genotype, groups_3152-5.
Supplementary Figure S7: Distribution of genomes containing a genotypes, groups_3152-6, -7, -22 and -37.
Supplementary Figure S8: Distribution of genomes containing a genotype, groups_3152-8.
Supplementary Figure S9: Distribution of genomes containing a genotype, groups_3152-9.
Supplementary Figure S10: Distribution of genomes containing a genotypes, groups_3152-10, -15, -27, -33, -34 and -35.
Supplementary Figure S11: Distribution of genomes containing a genotype, groups_3152-11.
Supplementary Figure S12: Distribution of genomes containing a genotypes, groups_3152-12 and -24.
Supplementary Figure S13: Distribution of genomes containing a genotypes, groups_3152-13, -20, -32 and -43.
Supplementary Figure S14: Distribution of genomes containing a genotypes, groups_3152-14, -17, -19 and -21.
Supplementary Figure S15: Distribution of genomes containing a genotypes, groups_3152-16, -23, -31, -38, -40 and -41.
Supplementary Figure S16: Distribution of genomes containing a genotypes, groups_3152-18 and -29.
Supplementary Figure S17: Distribution of genomes containing a genotypes, groups_3152-25, -26, -30 and -42.

Supplementary Figure S18: Distribution of genomes containing a genotypes, groups_3152-28, -36 and -39.

Supplementary Figure S1

#### A.One letter notation

| *Citrobacter* spp. | Code |
|---|---|
| *C. freundii* | D |
| *C. braakii* | B |
| *C. portucalensis* | P |
| *C. koseri* | K |
| *C. gillenii* | G |
| *C. amalonaticus* | L |
| *C. cronae* | C |
| *C. werkmanii* | W |
| *C. youngae* | Y |
| *C. europaeus* | E |
| *C. farmeri* | F |
| *C. meridianamericanus* | M |
| *C. pasteurii* | U |
| *C. rodentium* | R |
| *C. sedlakii* | S |
| *C. enshiensis* | N |
| *C. arsenatis* | A |
| *C. tructae* | T |

#### B. Coding rule in this study

#### C. Proposing coding rule

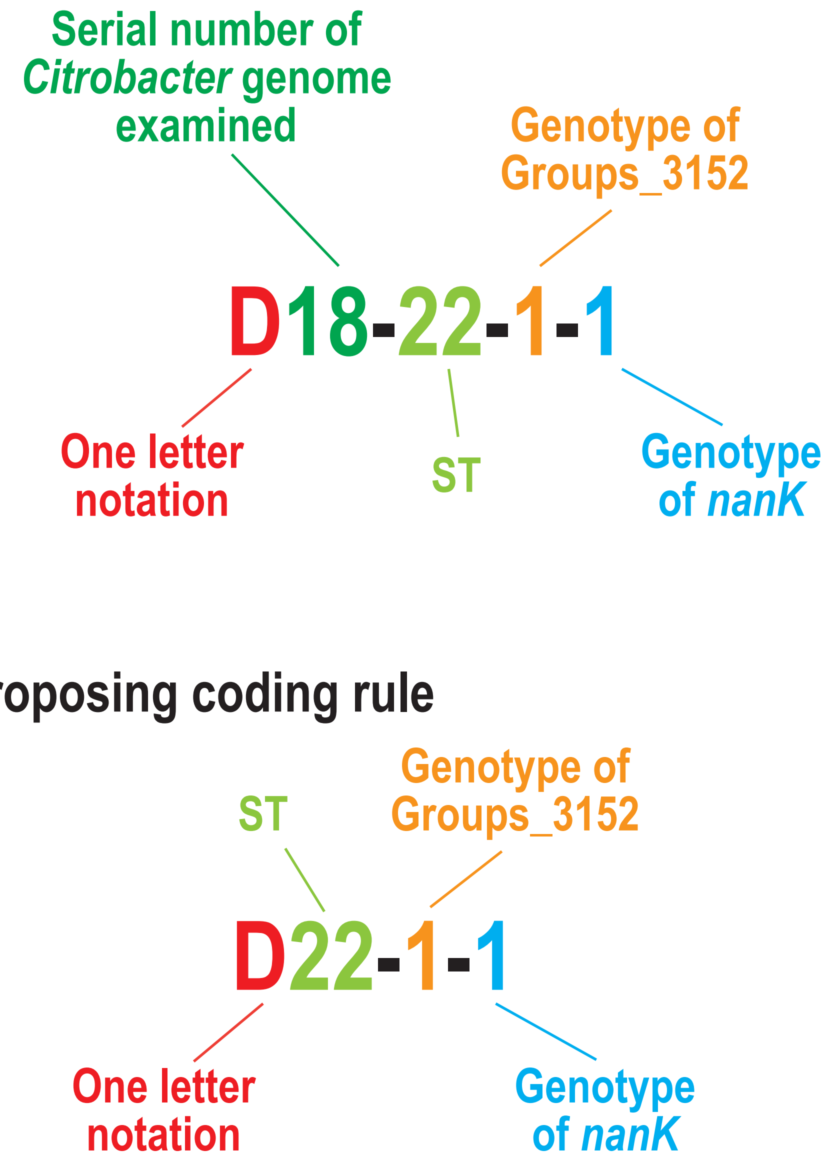

#### Supplementary Figure S2

**A**

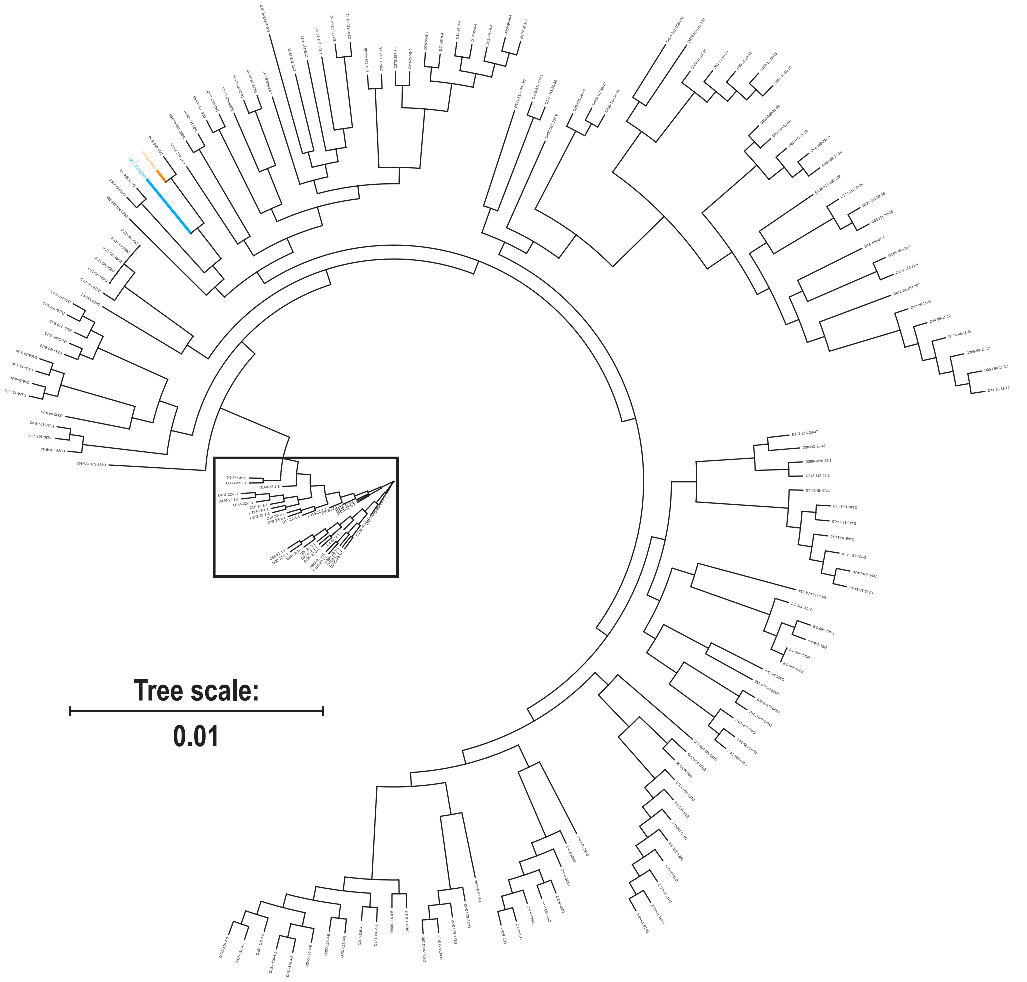

### Supplementary Figure S2

**B**

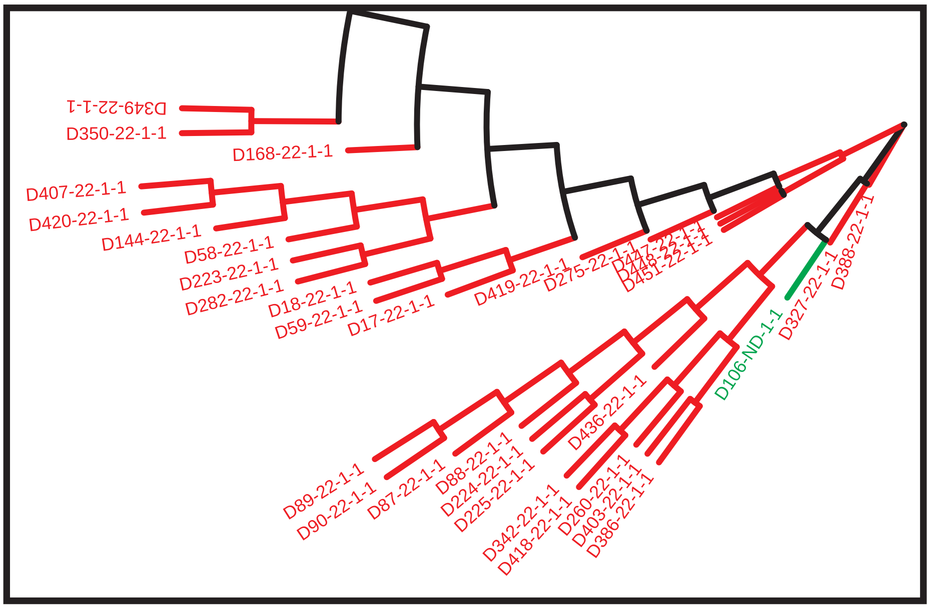


**C**

| | | | Genotypes of the top seven HND genes. | | | | | | |
|---|---|---|---|---|---|---|---|---|---|
| Coding | Accession # | ST | groups_3152 | *nanK* | *iprA* | *mipA* | *yehY* | *yhcH* | *ymdB* |
| D17-22-1-1 | CP011612.1 | 22 | 1 | 1 | 2 | 3 | 1 | 1 | 1 |
| D18-22-1-1 | CP011657.1 | 22 | 1 | 1 | 2 | 3 | 1 | 1 | 1 |
| D58-22-1-1 | CP036435.1 | 22 | 1 | 1 | 2 | 3 | 1 | 1 | 1 |
| D59-22-1-1 | CP037734.1 | 22 | 1 | 1 | 2 | 3 | 1 | 1 | 1 |
| D87-22-1-1 | CP047269.1 | 22 | 1 | 1 | 2 | 3 | 1 | 1 | 1 |
| D88-22-1-1 | CP047273.1 | 22 | 1 | 1 | 2 | 3 | 1 | 1 | 1 |
| D89-22-1-1 | CP047275.1 | 22 | 1 | 1 | 2 | 3 | 1 | 1 | 1 |
| D90-22-1-1 | CP047279.1 | 22 | 1 | 1 | 2 | 3 | 1 | 1 | 1 |
| D144-22-1-1 | CP056365.1 | 22 | 1 | 1 | 2 | 3 | 1 | 1 | 1 |
| D223-22-1-1 | CP071265.1 | 22 | 1 | 1 | 2 | 3 | 1 | 1 | 1 |
| D224-22-1-1 | CP071834.1 | 22 | 1 | 1 | 2 | 3 | 1 | 1 | 1 |
| D225-22-1-1 | CP071907.1 | 22 | 1 | 1 | 2 | 3 | 1 | 1 | 1 |
| D260-22-1-1 | CP086287.1 | 22 | 1 | 1 | 2 | 3 | 1 | 1 | 1 |
| D275-22-1-1 | CP098330.1 | 22 | 1 | 1 | 2 | 3 | 1 | 1 | 1 |
| D327-22-1-1 | CP110775.1 | 22 | 1 | 1 | 2 | 3 | 1 | 1 | 1 |
| D342-22-1-1 | CP117475.1 | 22 | 1 | 1 | 2 | 3 | 1 | 1 | 1 |
| D386-22-1-1 | CP137175.1 | 22 | 1 | 1 | 2 | 3 | 1 | 1 | 1 |
| D388-22-1-1 | CP137183.1 | 22 | 1 | 1 | 2 | 3 | 1 | 1 | 1 |
| D403-22-1-1 | CP141645.1 | 22 | 1 | 1 | 2 | 3 | 1 | 1 | 1 |
| D407-22-1-1 | CP145665.1 | 22 | 1 | 1 | 2 | 3 | 1 | 1 | 1 |
| D418-22-1-1 | CP162975.1 | 22 | 1 | 1 | 2 | 3 | 1 | 1 | 1 |
| D419-22-1-1 | CP162982.1 | 22 | 1 | 1 | 2 | 3 | 1 | 1 | 1 |
| D420-22-1-1 | CP163076.1 | 22 | 1 | 1 | 2 | 3 | 1 | 1 | 1 |
| D436-22-1-1 | LS992175.1 | 22 | 1 | 1 | 2 | 3 | 1 | 1 | 1 |
| D447-22-1-1 | OW849527.1 | 22 | 1 | 1 | 2 | 3 | 1 | 1 | 1 |
| D448-22-1-1 | OW967263.1 | 22 | 1 | 1 | 2 | 3 | 1 | 1 | 1 |
| D451-22-1-1 | OW969875.1 | 22 | 1 | 1 | 2 | 3 | 1 | 1 | 1 |
| D168-22-1-1 | CP056653.1 | 22 | 1 | 1 | 2 | 3 | 1 | 98 | 1 |
| D282-22-1-1 | CP099291.1 | 22 | 1 | 1 | 2 | 3 | 164 | 123 | 1 |
| D349-22-1-1 | CP124809.1 | 22 | 1 | 1 | 67 | 3 | 16 | 1 | 1 |
| D350-22-1-1 | CP124813.1 | 22 | 1 | 1 | 67 | 3 | 16 | 1 | 1 |
| D278-911-1-161 | CP099128.1 | 911 | 1 | 161 | 6 | 5 | 161 | 120 | 52 |
| D106-ND-1-1 | CP054294.1 | ND | 1 | 1 | 2 | 3 | 1 | 1 | 1 |
| D444-ND-1-2 | OW849099.1 | ND | 1 | 2 | 6 | 5 | 48 | 1 | 9 |

**Supplementary Figure S3**

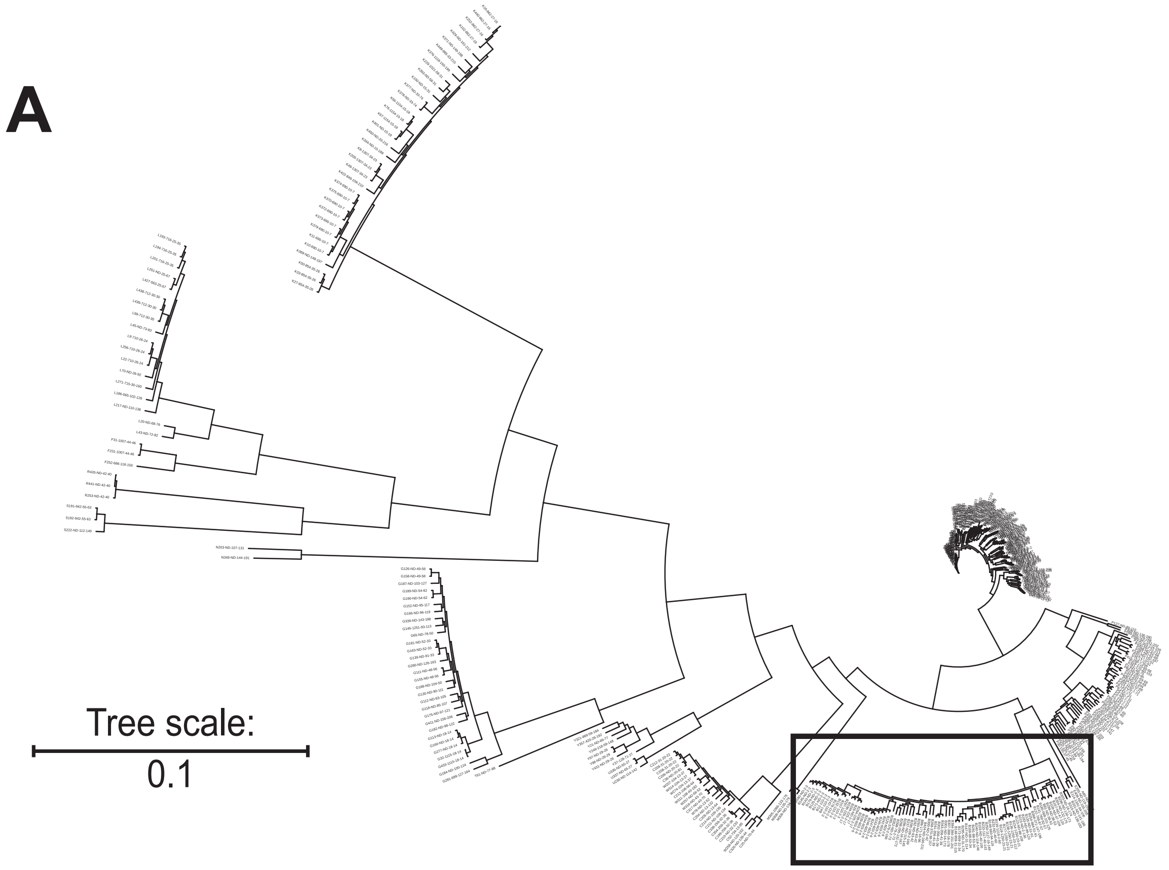

Supplementary Figure S3

**B**

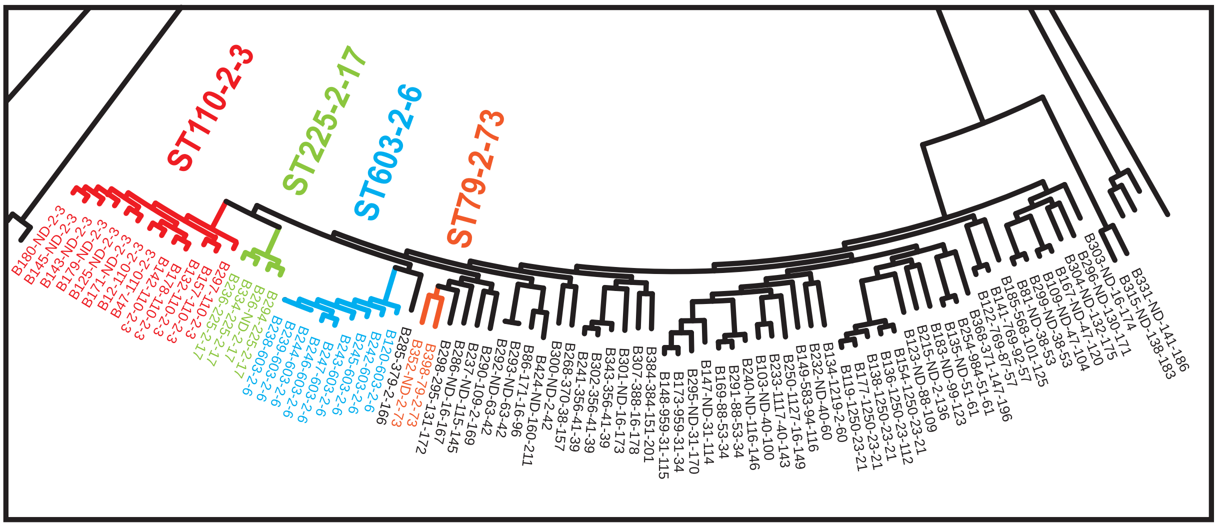


**C**

| | | | Genotypes of the top seven HND genes. | | | | | | |
|---|---|---|---|---|---|---|---|---|---|
| Coding # | Accession # | ST | groups_ 3152 | *nanK* | *iprA* | *mipA* | *yehY* | *yhcH* | *ymdB* |
| B12-110-2-3 | AP026382.1 | 110 | 2 | 3 | 4 | 4 | 2 | 3 | 3 |
| B47-110-2-3 | CP026235.1 | 110 | 2 | 3 | 4 | 4 | 2 | 3 | 3 |
| B132-110-2-3 | CP056251.1 | 110 | 2 | 3 | 4 | 4 | 2 | 3 | 3 |
| B142-110-2-3 | CP056350.1 | 110 | 2 | 3 | 4 | 4 | 2 | 3 | 3 |
| B157-110-2-3 | CP056546.1 | 110 | 2 | 3 | 4 | 4 | 2 | 3 | 3 |
| B178-110-2-3 | CP056888.1 | 110 | 2 | 3 | 4 | 4 | 2 | 3 | 3 |
| B297-110-2-3 | CP099382.1 | 110 | 2 | 3 | 4 | 4 | 2 | 3 | 3 |
| B125-ND-2-3 | CP056219.1 | ND | 2 | 3 | 4 | 4 | 2 | 3 | 3 |
| B143-ND-2-3 | CP056361.1 | ND | 2 | 3 | 4 | 4 | 2 | 3 | 3 |
| B145-ND-2-3 | CP056381.1 | ND | 2 | 3 | 4 | 4 | 2 | 3 | 3 |
| B171-ND-2-3 | CP056822.1 | ND | 2 | 3 | 4 | 4 | 2 | 3 | 3 |
| B179-ND-2-3 | CP056896.1 | ND | 2 | 3 | 4 | 4 | 2 | 3 | 3 |
| B180-ND-2-3 | CP056899.1 | ND | 2 | 3 | 4 | 4 | 2 | 3 | 3 |
| B120-603-2-6 | CP056180.1 | 603 | 2 | 6 | 7 | 8 | 6 | 10 | 4 |
| B238-603-2-6 | CP078595.1 | 603 | 2 | 6 | 7 | 8 | 6 | 10 | 4 |
| B239-603-2-6 | CP078596.1 | 603 | 2 | 6 | 7 | 8 | 6 | 10 | 4 |
| B242-603-2-6 | CP078599.1 | 603 | 2 | 6 | 7 | 8 | 6 | 10 | 4 |
| B243-603-2-6 | CP078600.1 | 603 | 2 | 6 | 7 | 8 | 6 | 10 | 4 |
| B244-603-2-6 | CP078601.1 | 603 | 2 | 6 | 7 | 8 | 6 | 10 | 4 |
| B245-603-2-6 | CP078602.1 | 603 | 2 | 6 | 7 | 8 | 6 | 10 | 4 |
| B246-603-2-6 | CP078603.1 | 603 | 2 | 6 | 7 | 8 | 6 | 10 | 4 |
| B247-603-2-6 | CP078604.1 | 603 | 2 | 6 | 7 | 8 | 6 | 10 | 4 |
| B236-225-2-17 | CP078593.1 | 225 | 2 | 17 | 13 | 16 | 23 | 23 | 28 |
| B294-225-2-17 | CP099378.1 | 225 | 2 | 17 | 13 | 16 | 23 | 23 | 28 |
| B334-225-2-17 | CP114801.1 | 225 | 2 | 17 | 13 | 16 | 23 | 23 | 28 |
| B26-ND-2-17 | CP022049.2 | ND | 2 | 17 | 13 | 16 | 23 | 23 | 28 |
| B300-ND-2-42 | CP099386.1 | ND | 2 | 42 | 139 | 16 | 170 | 125 | 132 |
| B134-1219-2-60 | CP056267.1 | 1219 | 2 | 60 | 100 | 90 | 115 | 95 | 14 |
| B398-79-2-73 | CP138570.1 | 79 | 2 | 73 | 7 | 61 | 202 | 43 | 72 |
| B352-ND-2-73 | CP126329.1 | ND | 2 | 73 | 17 | 4 | 2 | 43 | 146 |
| B215-ND-2-136 | CP069788.1 | ND | 2 | 136 | 120 | 11 | 138 | 61 | 3 |
| B285-379-2-166 | CP099364.1 | 379 | 2 | 166 | 63 | 51 | 2 | 61 | 72 |
| B290-109-2-169 | CP099374.1 | 109 | 2 | 169 | 7 | 124 | 168 | 57 | 3 |

#### Supplementary Figure S4

A

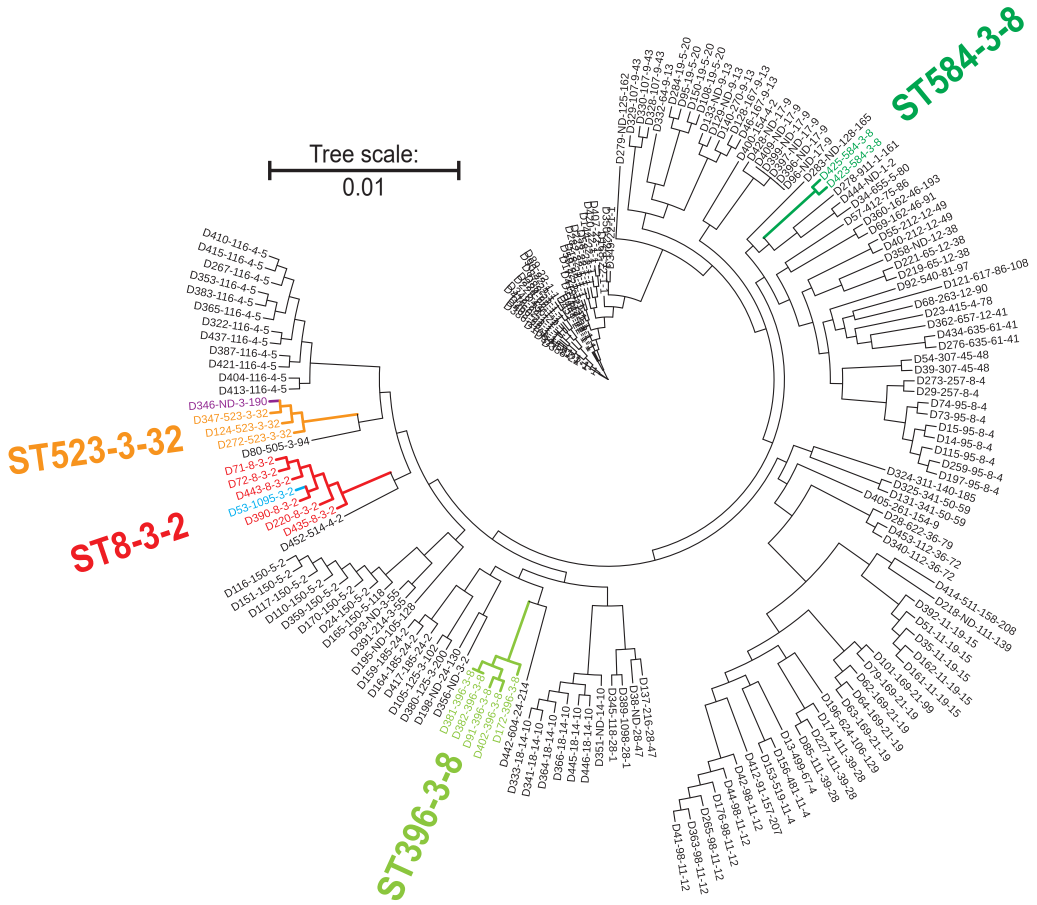


B

| | | | Genotypes of the top seven HND genes. | | | | | | |
|---|---|---|---|---|---|---|---|---|---|
| Coding # | Accession # | ST | groups_3152 | *nanK* | *iprA* | *mipA* | *yehY* | *yhcH* | *ymdB* |
| D71-8-3-2 | CP042478.1 | 8 | 3 | 2 | 1 | 1 | 11 | 1 | 10 |
| D72-8-3-2 | CP042517.1 | 8 | 3 | 2 | 1 | 1 | 11 | 1 | 10 |
| D220-8-3-2 | CP070549.1 | 8 | 3 | 2 | 1 | 1 | 11 | 1 | 10 |
| D390-8-3-2 | CP137201.1 | 8 | 3 | 2 | 1 | 1 | 11 | 1 | 10 |
| D435-8-3-2 | LR890181.1 | 8 | 3 | 2 | 1 | 1 | 215 | 1 | 10 |
| D443-8-3-2 | OW849082.1 | 8 | 3 | 2 | 1 | 1 | 11 | 1 | 10 |
| D53-1095-3-2 | CP032184.1 | 1095 | 3 | 2 | 1 | 1 | 11 | 1 | 10 |
| D356-ND-3-2 | CP126623.1 | ND | 3 | 2 | 36 | 2 | 76 | 1 | 147 |
| D91-396-3-8 | CP047307.1 | 396 | 3 | 8 | 18 | 5 | 17 | 2 | 24 |
| D172-396-3-8 | CP056827.1 | 396 | 3 | 8 | 18 | 5 | 17 | 2 | 24 |
| D381-396-3-8 | CP137123.1 | 396 | 3 | 8 | 18 | 5 | 17 | 2 | 24 |
| D382-396-3-8 | CP137128.1 | 396 | 3 | 8 | 18 | 5 | 17 | 2 | 24 |
| D402-396-3-8 | CP140972.1 | 396 | 3 | 8 | 18 | 5 | 17 | 2 | 24 |
| D423-584-3-8 | CP167050.1 | 584 | 3 | 8 | 71 | 2 | 84 | 26 | 48 |
| D425-584-3-8 | CP167107.1 | 584 | 3 | 8 | 71 | 2 | 84 | 26 | 48 |
| D124-523-3-32 | CP056208.1 | 523 | 3 | 32 | 9 | 1 | 3 | 35 | 9 |
| D272-523-3-32 | CP096921.1 | 523 | 3 | 32 | 9 | 1 | 3 | 35 | 9 |
| D347-523-3-32 | CP119165.1 | 523 | 3 | 32 | 9 | 1 | 3 | 35 | 9 |
| D346-ND-3-190 | CP119053.1 | ND | 3 | 190 | 9 | 1 | 187 | 134 | 9 |
| D391-214-3-55 | CP137203.1 | 214 | 3 | 55 | 1 | 6 | 3 | 51 | 7 |
| D93-ND-3-55 | CP048382.1 | ND | 3 | 55 | 1 | 6 | 105 | 51 | 2 |
| D80-505-3-94 | CP045726.1 | 505 | 3 | 94 | 91 | 2 | 16 | 14 | 93 |
| D105-125-3-102 | CP054278.1 | 125 | 3 | 102 | 93 | 46 | 57 | 2 | 7 |
| D380-125-3-200 | CP137117.1 | 125 | 3 | 200 | 157 | 46 | 57 | 2 | 7 |

#### Supplementary Figure S5

**A**

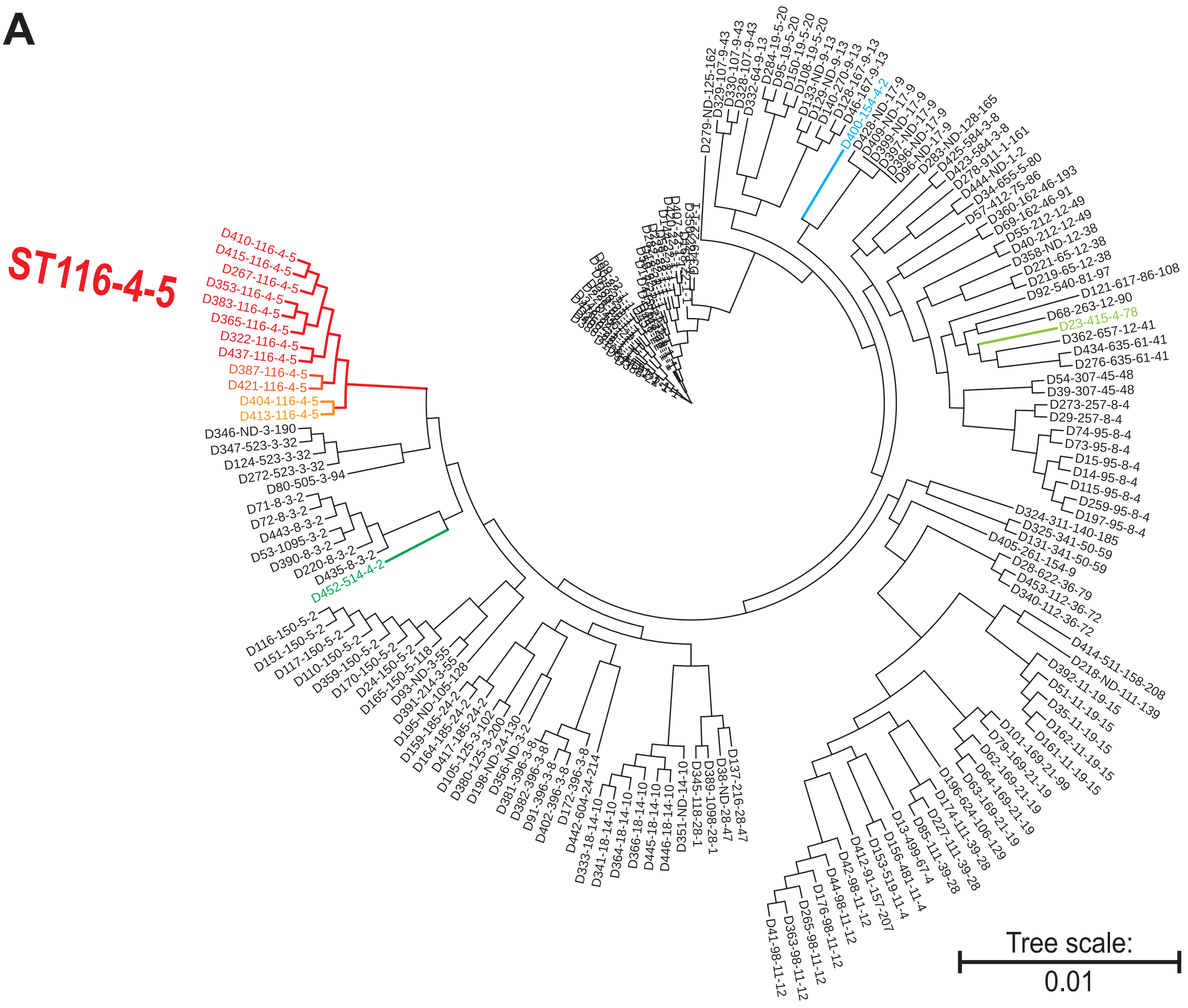


**B**

| Coding # | Accession # | ST | Genotypes of the top seven HND genes. | | | | | | |
|---|---|---|---|---|---|---|---|---|---|
| | | | groups_3152 | *nanK* | *iprA* | *mipA* | *yehY* | *yhcH* | *ymdB* |
| D267-116-4-5 | CP092493.1 | 116 | 4 | 5 | 5 | 2 | 4 | 6 | 2 |
| D322-116-4-5 | CP103365.1 | 116 | 4 | 5 | 5 | 2 | 4 | 6 | 2 |
| D353-116-4-5 | CP126536.1 | 116 | 4 | 5 | 5 | 2 | 4 | 6 | 2 |
| D365-116-4-5 | CP135622.1 | 116 | 4 | 5 | 5 | 2 | 4 | 6 | 2 |
| D383-116-4-5 | CP137133.1 | 116 | 4 | 5 | 5 | 2 | 4 | 6 | 2 |
| D410-116-4-5 | CP150622.1 | 116 | 4 | 5 | 5 | 2 | 4 | 6 | 2 |
| D415-116-4-5 | CP155122.1 | 116 | 4 | 5 | 5 | 2 | 4 | 6 | 2 |
| D437-116-4-5 | LS992183.1 | 116 | 4 | 5 | 5 | 2 | 4 | 6 | 2 |
| D387-116-4-5 | CP137179.1 | 116 | 4 | 5 | 5 | 2 | 4 | 6 | 78 |
| D421-116-4-5 | CP165768.1 | 116 | 4 | 5 | 5 | 2 | 4 | 6 | 78 |
| D404-116-4-5 | CP142904.1 | 116 | 4 | 5 | 5 | 2 | 16 | 6 | 2 |
| D413-116-4-5 | CP151860.1 | 116 | 4 | 5 | 5 | 2 | 16 | 6 | 2 |
| D400-154-4-2 | CP139850.1 | 154 | 4 | 2 | 44 | 146 | 10 | 142 | 9 |
| D23-415-4-78 | CP016762.1 | 415 | 4 | 78 | 73 | 70 | 88 | 2 | 82 |
| D452-514-4-2 | OW969904.1 | 514 | 4 | 2 | 1 | 6 | 49 | 1 | 164 |

#### Supplementary Figure S6

**A**

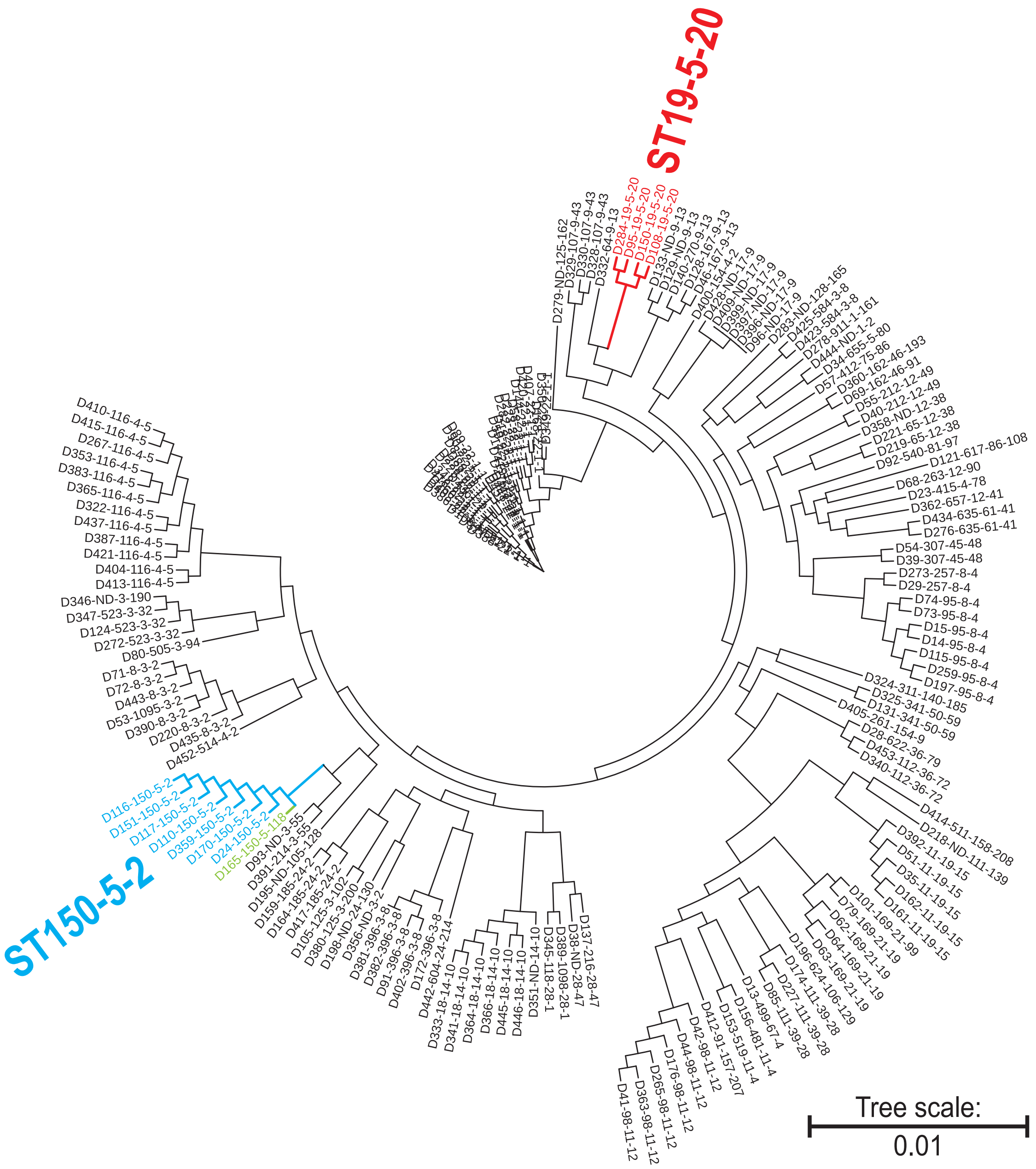


**B**

| | | | Genotypes of the top seven HND genes. | | | | | | |
|---|---|---|---|---|---|---|---|---|---|
| Coding # | Accession # | ST | groups_ 3152 | *nanK* | *iprA* | *mipA* | *yehY* | *yhcH* | *ymdB* |
| D95-19-5-20 | CP048416.1 | 19 | 5 | 20 | 8 | 45 | 18 | 2 | 1 |
| D150-19-5-20 | CP056451.1 | 19 | 5 | 20 | 8 | 45 | 18 | 2 | 1 |
| D108-19-5-20 | CP055247.1 | 19 | 5 | 20 | 8 | 85 | 18 | 2 | 1 |
| D284-19-5-20 | CP099303.1 | 19 | 5 | 20 | 8 | 122 | 18 | 2 | 1 |
| D24-150-5-2 | CP016952.1 | 150 | 5 | 2 | 12 | 2 | 22 | 1 | 2 |
| D170-150-5-2 | CP056809.1 | 150 | 5 | 2 | 12 | 2 | 22 | 1 | 2 |
| D359-150-5-2 | CP133060.1 | 150 | 5 | 2 | 12 | 2 | 22 | 1 | 2 |
| D110-150-5-2 | CP055421.1 | 150 | 5 | 2 | 12 | 2 | 25 | 1 | 2 |
| D116-150-5-2 | CP055582.1 | 150 | 5 | 2 | 12 | 2 | 25 | 1 | 2 |
| D117-150-5-2 | CP055588.1 | 150 | 5 | 2 | 12 | 2 | 25 | 1 | 2 |
| D151-150-5-2 | CP056466.1 | 150 | 5 | 2 | 12 | 2 | 25 | 1 | 2 |
| D165-150-5-118 | CP056635.1 | 150 | 5 | 118 | 103 | 2 | 22 | 1 | 2 |
| D34-655-5-80 | CP024672.1 | 655 | 5 | 80 | 6 | 5 | 48 | 74 | 9 |

#### Supplementary Figure S7

**A**

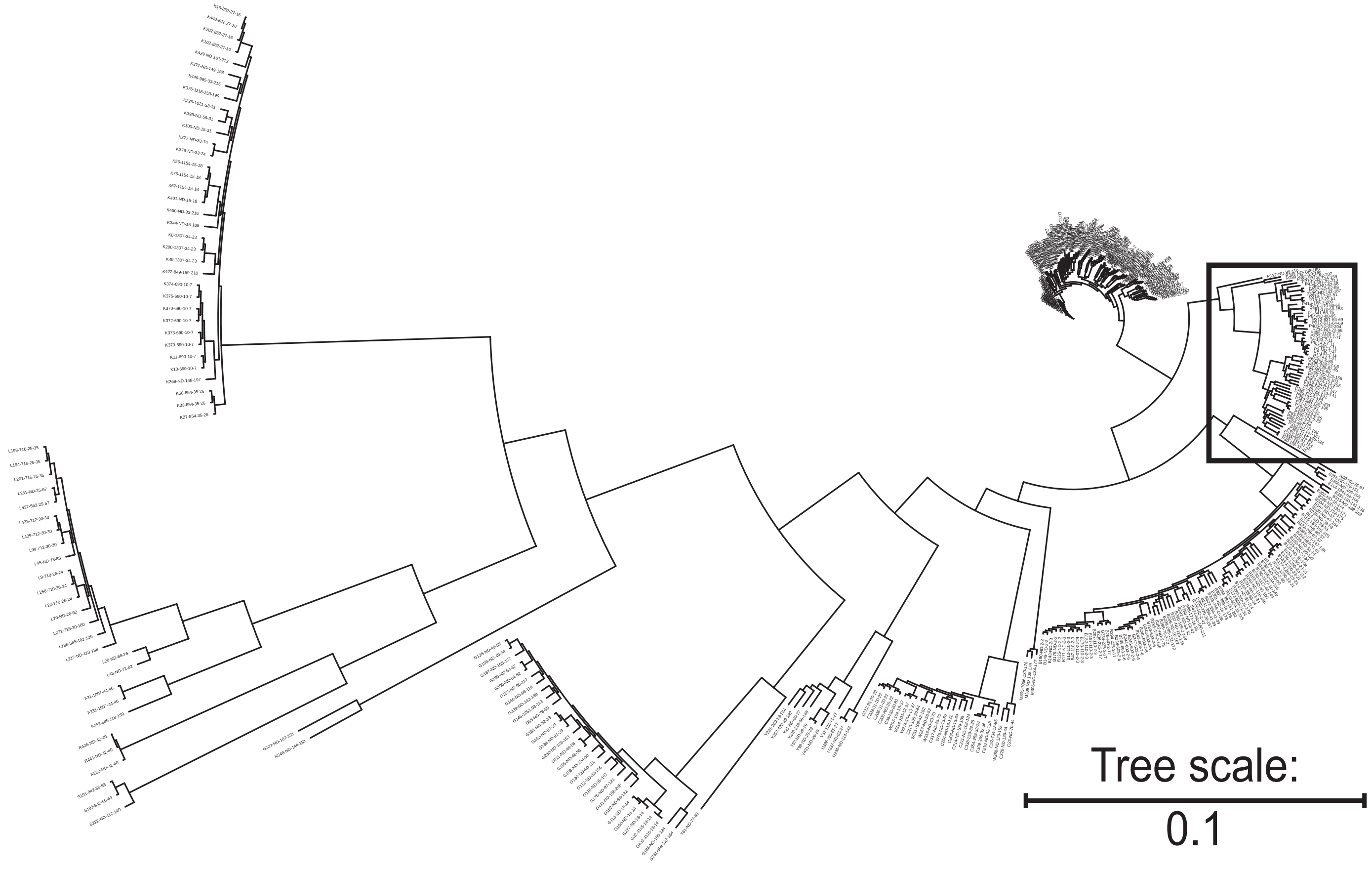

#### Supplementary Figure S7

**B**

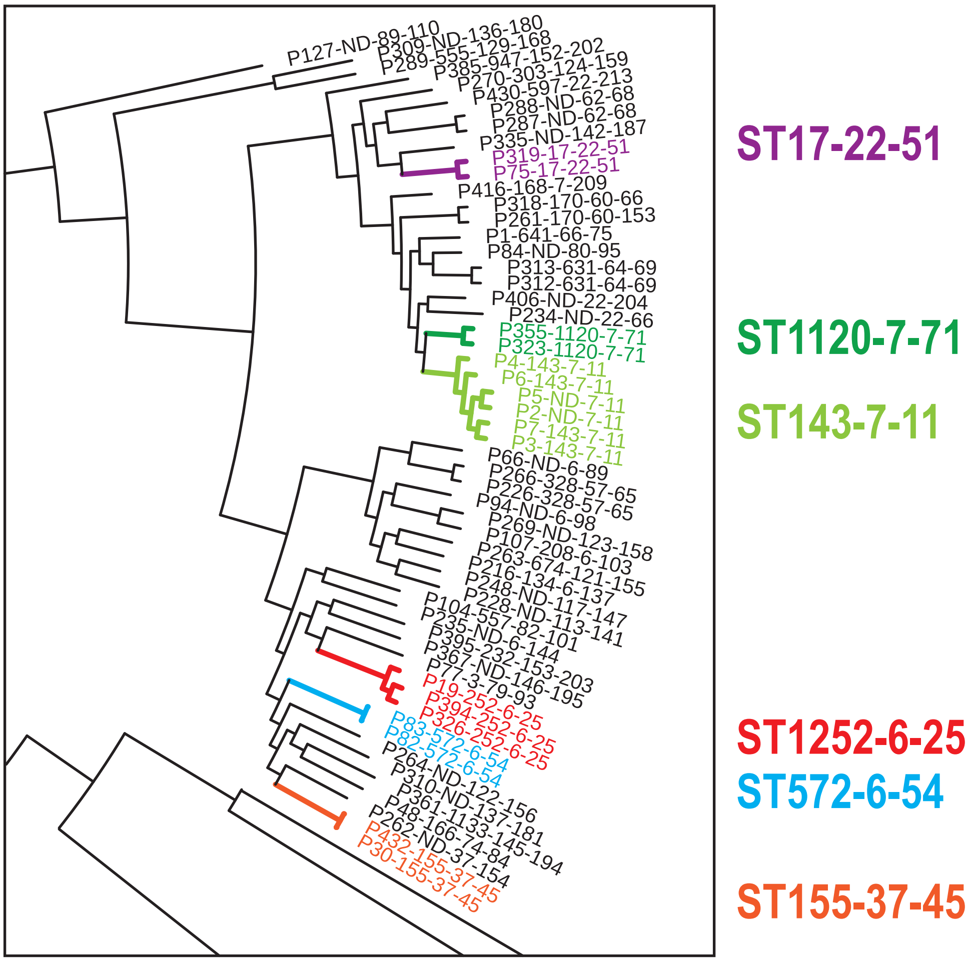


**C**

| | | | Genotypes of the top seven HND genes. | | | | | | |
|---|---|---|---|---|---|---|---|---|---|
| Coding # | Accession # | ST | groups_3152 | *nanK* | *iprA* | *mipA* | *yehY* | *yhcH* | *ymdB* |
| P216-134-6-137 | CP069801.1 | 134 | 6 | 137 | 25 | 101 | 139 | 18 | 116 |
| P107-208-6-103 | CP055089.1 | 208 | 6 | 103 | 29 | 47 | 58 | 18 | 58 |
| P19-252-6-25 | CP012554.1 | 252 | 6 | 25 | 23 | 18 | 28 | 22 | 36 |
| P326-252-6-25 | CP109744.1 | 252 | 6 | 25 | 23 | 18 | 28 | 22 | 36 |
| P394-252-6-25 | CP137478.1 | 252 | 6 | 25 | 23 | 18 | 28 | 22 | 36 |
| P82-572-6-54 | CP045837.1 | 572 | 6 | 54 | 29 | 44 | 54 | 12 | 41 |
| P83-572-6-54 | CP045840.1 | 572 | 6 | 54 | 29 | 44 | 54 | 12 | 41 |
| P66-ND-6-89 | CP039327.1 | ND | 6 | 89 | 88 | 80 | 30 | 81 | 92 |
| P94-ND-6-98 | CP048388.1 | ND | 6 | 98 | 25 | 83 | 106 | 18 | 96 |
| P235-ND-6-144 | CP078551.1 | ND | 6 | 144 | 16 | 108 | 147 | 12 | 41 |
| P3-143-7-11 | AP022394.1 | 143 | 7 | 11 | 3 | 13 | 8 | 8 | 5 |
| P4-143-7-11 | AP022399.1 | 143 | 7 | 11 | 3 | 13 | 8 | 8 | 5 |
| P6-143-7-11 | AP022494.1 | 143 | 7 | 11 | 3 | 13 | 8 | 8 | 5 |
| P7-143-7-11 | AP022513.1 | 143 | 7 | 11 | 3 | 13 | 8 | 8 | 5 |
| P2-ND-7-11 | AP022389.1 | ND | 7 | 11 | 3 | 13 | 8 | 8 | 5 |
| P5-ND-7-11 | AP022486.1 | ND | 7 | 11 | 3 | 13 | 8 | 8 | 5 |
| P416-168-7-209 | CP159117.1 | 168 | 7 | 209 | 164 | 7 | 209 | 7 | 54 |
| P323-1120-7-71 | CP104921.1 | 1120 | 7 | 71 | 3 | 7 | 81 | 70 | 5 |
| P355-1120-7-71 | CP126611.1 | 1120 | 7 | 71 | 3 | 7 | 81 | 70 | 5 |
| P75-17-22-51 | CP043009.1 | 17 | 22 | 51 | 47 | 28 | 31 | 7 | 31 |
| P319-17-22-51 | CP101100.1 | 17 | 22 | 51 | 47 | 28 | 31 | 7 | 31 |
| P430-597-22-213 | LR134214.1 | 597 | 22 | 213 | 70 | 151 | 213 | 147 | 31 |
| P234-ND-22-66 | CP078550.1 | ND | 22 | 66 | 3 | 7 | 146 | 8 | 49 |
| P406-ND-22-204 | CP145154.1 | ND | 22 | 204 | 3 | 7 | 204 | 7 | 159 |
| P30-155-37-45 | CP022311.1 | 155 | 37 | 45 | 25 | 39 | 45 | 12 | 55 |
| P432-155-37-45 | LR698971.1 | 155 | 37 | 45 | 25 | 39 | 45 | 12 | 55 |
| P262-ND-37-154 | CP089437.1 | ND | 37 | 154 | 23 | 115 | 155 | 12 | 127 |

#### Supplementary Figure S8

**A**

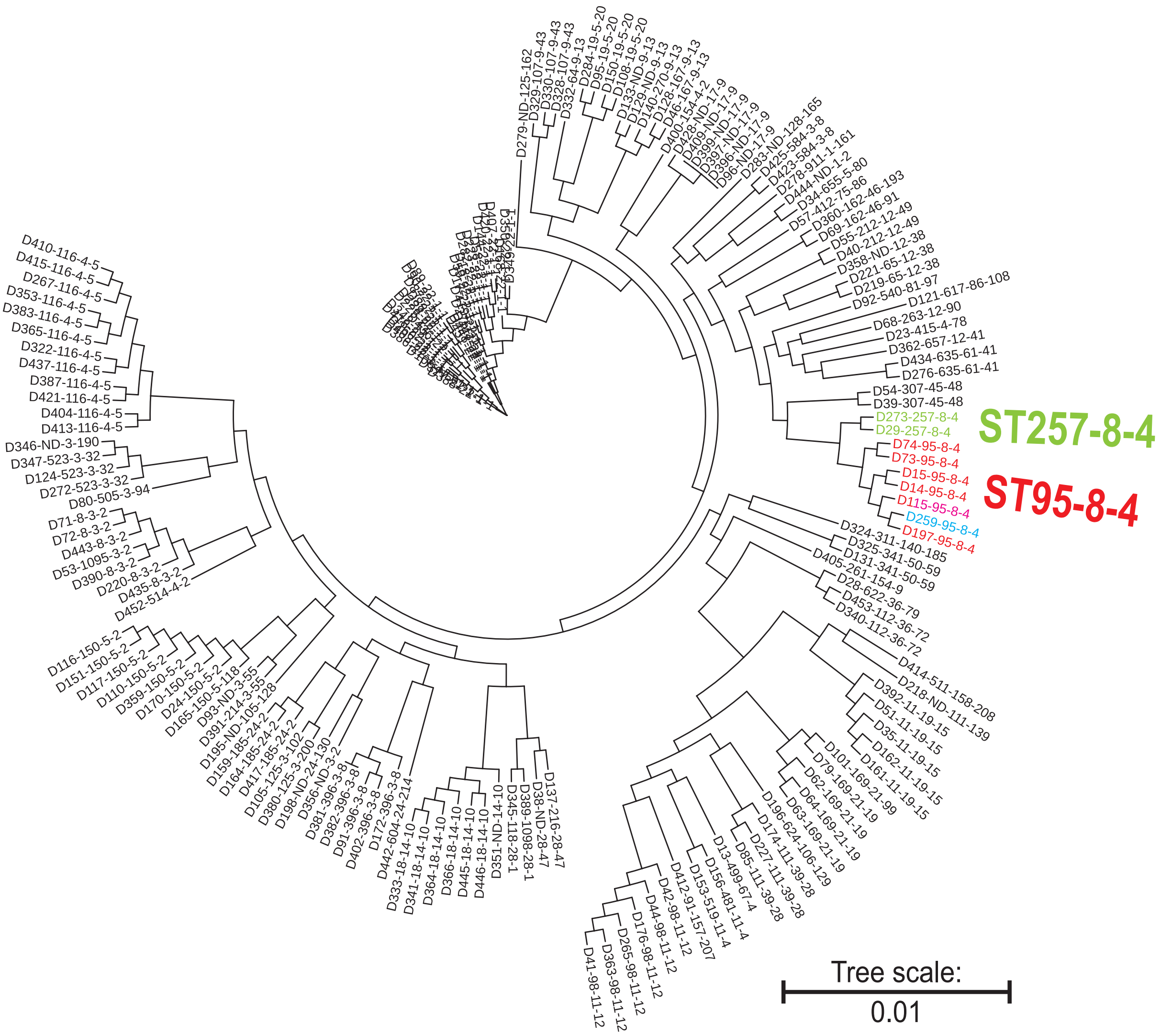


**B**

| | | | Genotypes of the top seven HND genes. | | | | | | |
|---|---|---|---|---|---|---|---|---|---|
| Coding # | Accession # | ST | groups_3152 | *nanK* | *iprA* | *mipA* | *yehY* | *yhcH* | *ymdB* |
| D14-95-8-4 | AP028314.1 | 95 | 8 | 4 | 1 | 10 | 5 | 4 | 8 |
| D15-95-8-4 | AP028317.1 | 95 | 8 | 4 | 1 | 10 | 5 | 4 | 8 |
| D73-95-8-4 | CP042524.1 | 95 | 8 | 4 | 1 | 10 | 5 | 4 | 8 |
| D74-95-8-4 | CP042534.1 | 95 | 8 | 4 | 1 | 10 | 5 | 4 | 8 |
| D115-95-8-4 | CP055564.1 | 95 | 8 | 4 | 1 | 10 | 5 | 4 | 8 |
| D197-95-8-4 | CP059427.1 | 95 | 8 | 4 | 1 | 10 | 5 | 4 | 8 |
| D259-95-8-4 | CP085726.1 | 95 | 8 | 4 | 1 | 114 | 5 | 4 | 8 |
| D29-257-8-4 | CP022273.1 | 257 | 8 | 4 | 1 | 10 | 5 | 4 | 20 |
| D273-257-8-4 | CP097107.1 | 257 | 8 | 4 | 1 | 10 | 5 | 4 | 20 |

#### Supplementary Figure S9

**A**

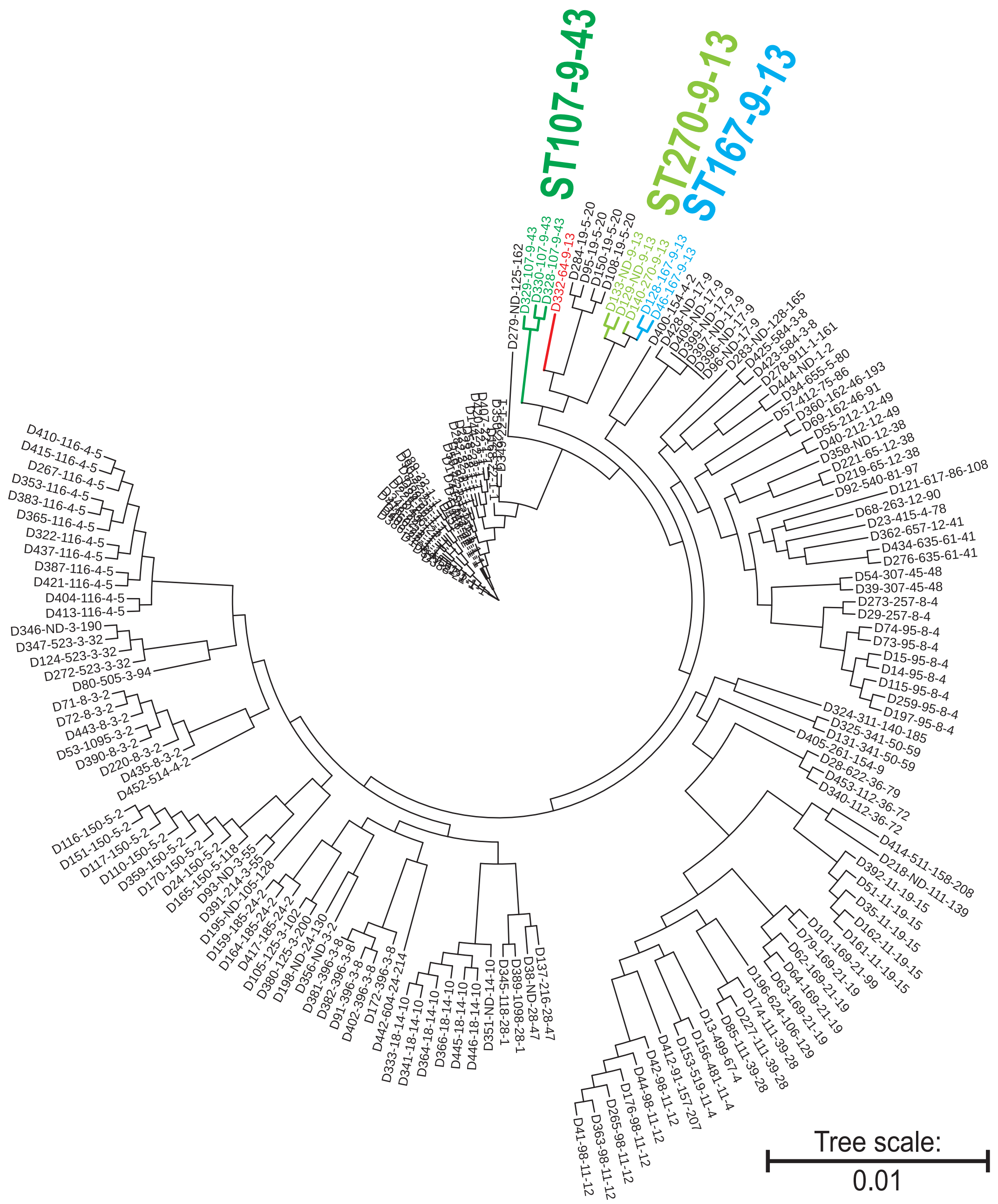


**B**

| | | | Genotypes of the seven HND genes. | | | | | | |
|---|---|---|---|---|---|---|---|---|---|
| Coding # | Accession # | ST | groups_ 3152 | *nanK* | *iprA* | *mipA* | *yehY* | *yhcH* | *ymdB* |
| D332-64-9-13 | CP113784.1 | 64 | 9 | 13 | 8 | 6 | 18 | 1 | 1 |
| D46-167-9-13 | CP026231.1 | 167 | 9 | 13 | 8 | 1 | 10 | 1 | 23 |
| D128-167-9-13 | CP056235.1 | 167 | 9 | 13 | 8 | 1 | 10 | 1 | 23 |
| D140-270-9-13 | CP056336.1 | 270 | 9 | 13 | 8 | 1 | 10 | 1 | 23 |
| D129-ND-9-13 | CP056238.1 | ND | 9 | 13 | 8 | 49 | 10 | 1 | 23 |
| D133-ND-9-13 | CP056256.1 | ND | 9 | 13 | 8 | 49 | 10 | 1 | 23 |
| D328-107-9-43 | CP110894.1 | 107 | 9 | 43 | 1 | 2 | 42 | 4 | 53 |
| D329-107-9-43 | CP110900.1 | 107 | 9 | 43 | 1 | 2 | 42 | 4 | 53 |
| D330-107-9-43 | CP110914.1 | 107 | 9 | 43 | 1 | 2 | 42 | 4 | 53 |

**Supplementary Figure S10**

**A**

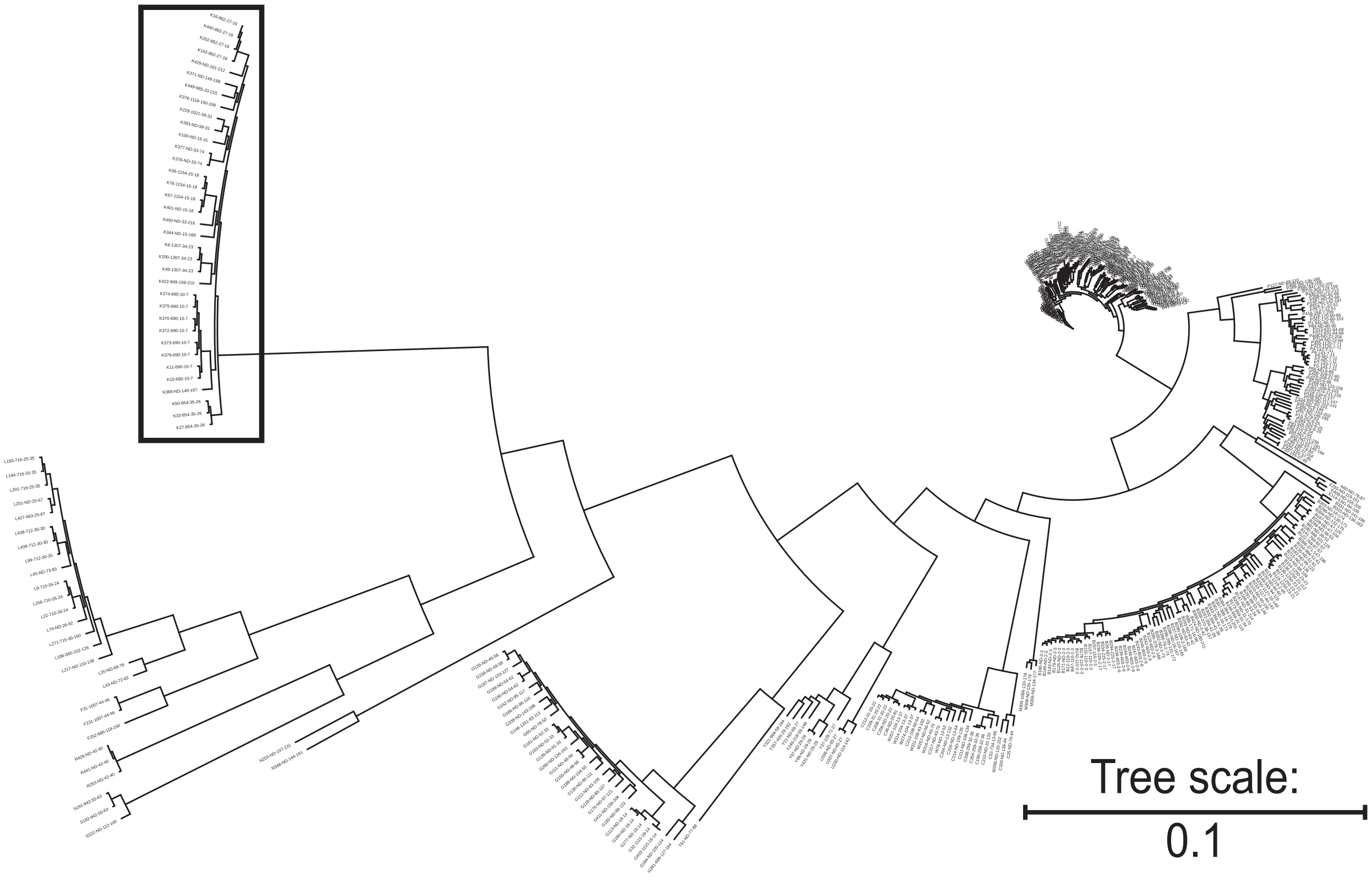

#### Supplementary Figure S10

**B**

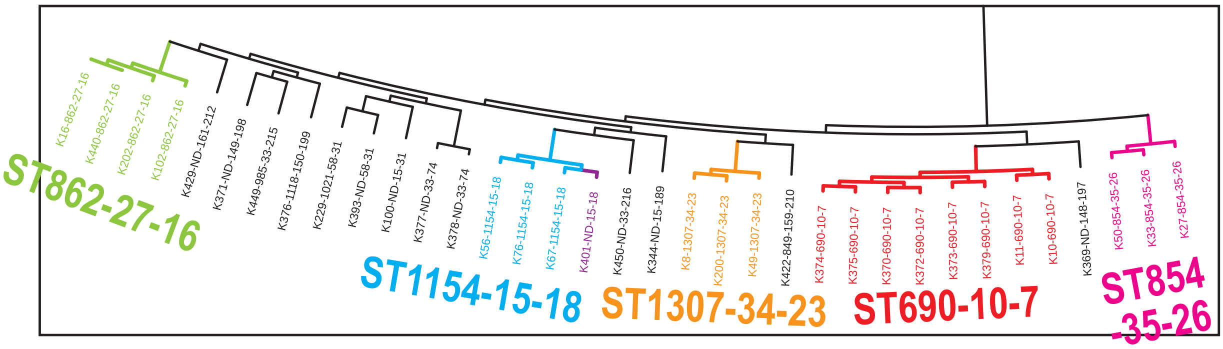


**C**

| | | | Genotypes of the top seven HND genes. | | | | | | |
|---|---|---|---|---|---|---|---|---|---|
| Coding | Accession # | ST | groups_ 3152 | *nanK* | *iprA* | *mipA* | *yehY* | *yhcH* | *ymdB* |
| K10-690-10-7 | AP025640.1 | 690 | 10 | 7 | 11 | 9 | 7 | 11 | 6 |
| K11-690-10-7 | AP025653.1 | 690 | 10 | 7 | 11 | 9 | 7 | 11 | 6 |
| K370-690-10-7 | CP136808.1 | 690 | 10 | 7 | 11 | 9 | 7 | 11 | 6 |
| K372-690-10-7 | CP136813.1 | 690 | 10 | 7 | 11 | 9 | 7 | 11 | 6 |
| K373-690-10-7 | CP136816.1 | 690 | 10 | 7 | 11 | 9 | 7 | 11 | 6 |
| K374-690-10-7 | CP136819.1 | 690 | 10 | 7 | 11 | 9 | 7 | 11 | 6 |
| K375-690-10-7 | CP136821.1 | 690 | 10 | 7 | 11 | 9 | 7 | 11 | 6 |
| K379-690-10-7 | CP137005.1 | 690 | 10 | 7 | 11 | 9 | 7 | 11 | 6 |
| K56-1154-15-18 | CP033780.1 | 1154 | 15 | 18 | 26 | 23 | 24 | 20 | 30 |
| K67-1154-15-18 | CP040234.1 | 1154 | 15 | 18 | 26 | 23 | 24 | 20 | 30 |
| K76-1154-15-18 | CP044097.1 | 1154 | 15 | 18 | 26 | 23 | 24 | 20 | 30 |
| K401-ND-15-18 | CP139989.1 | ND | 15 | 18 | 26 | 23 | 24 | 20 | 30 |
| K100-ND-15-31 | CP050078.1 | ND | 15 | 31 | 48 | 84 | 34 | 85 | 32 |
| K344-ND-15-189 | CP118927.1 | ND | 15 | 189 | 24 | 135 | 186 | 71 | 77 |
| K16-862-27-16 | CP000822.1 | 862 | 27 | 16 | 22 | 22 | 21 | 21 | 26 |
| K102-862-27-16 | CP052059.1 | 862 | 27 | 16 | 22 | 22 | 21 | 21 | 26 |
| K202-862-27-16 | CP066089.1 | 862 | 27 | 16 | 22 | 22 | 21 | 21 | 26 |
| K440-862-27-16 | NC_009792.1 | 862 | 27 | 16 | 22 | 22 | 21 | 21 | 26 |
| K377-ND-33-74 | CP136828.1 | ND | 33 | 74 | 69 | 37 | 83 | 19 | 32 |
| K378-ND-33-74 | CP136832.1 | ND | 33 | 74 | 69 | 37 | 83 | 19 | 32 |
| K449-985-33-215 | OW969691.1 | 985 | 33 | 215 | 169 | 37 | 216 | 149 | 79 |
| K450-ND-33-216 | OW969711.1 | ND | 33 | 216 | 170 | 153 | 217 | 20 | 77 |
| K8-1307-34-23 | AP023452.1 | 1307 | 34 | 23 | 31 | 14 | 20 | 30 | 34 |
| K49-1307-34-23 | CP026697.1 | 1307 | 34 | 23 | 31 | 14 | 20 | 30 | 34 |
| K200-1307-34-23 | CP060484.1 | 1307 | 34 | 23 | 31 | 14 | 20 | 30 | 34 |
| K27-854-35-26 | CP022073.2 | 854 | 35 | 26 | 24 | 14 | 29 | 19 | 37 |
| K33-854-35-26 | CP023527.1 | 854 | 35 | 26 | 24 | 14 | 29 | 19 | 37 |
| K50-854-35-26 | CP026709.1 | 854 | 35 | 26 | 24 | 14 | 29 | 19 | 37 |

#### Supplementary Figure S11

**A**

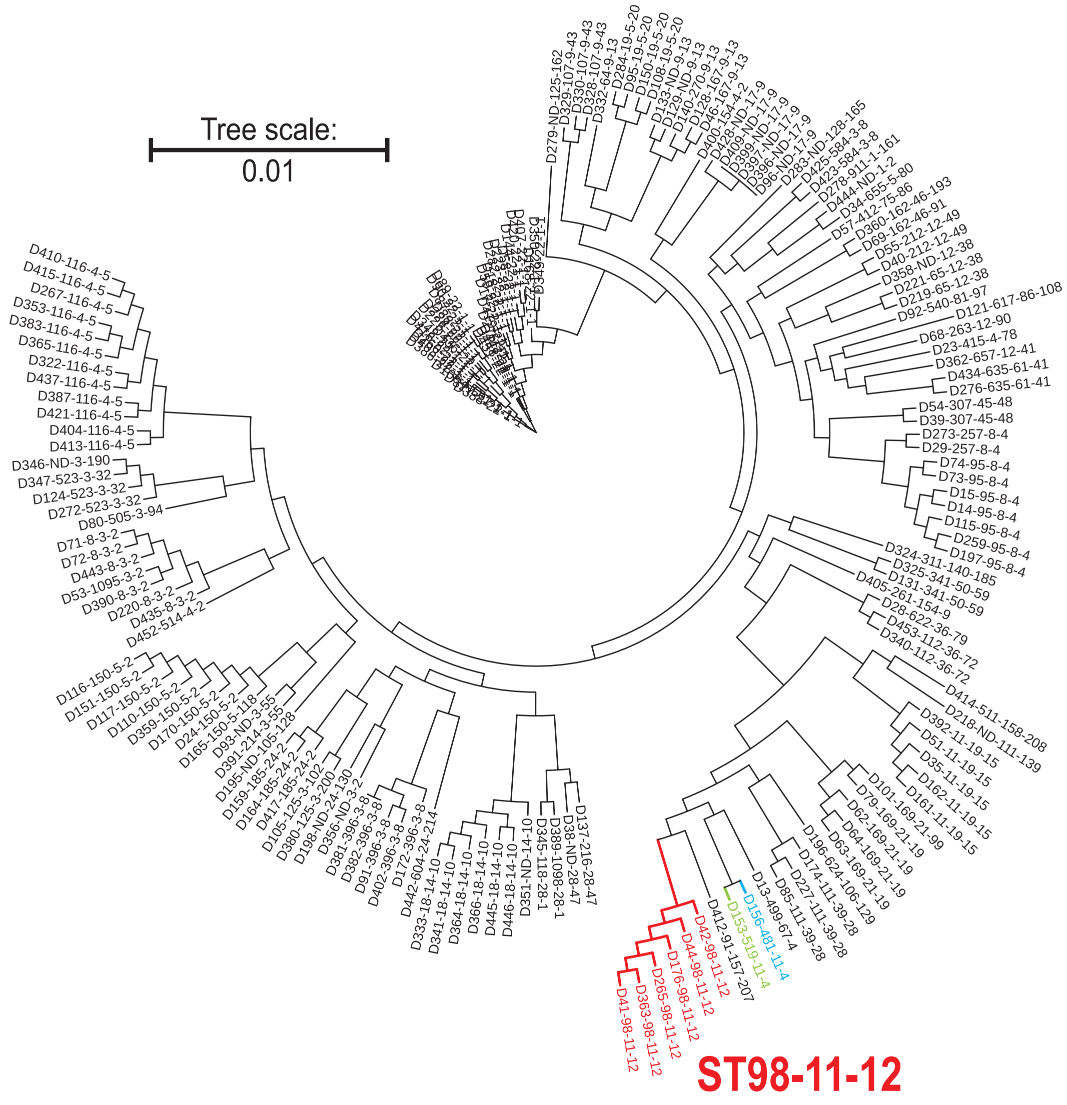


**B**

| | | | Genotypes of the top seven HND genes. | | | | | | |
|---|---|---|---|---|---|---|---|---|---|
| Coding | Accession # | ST | groups_3152 | *nanK* | *iprA* | *mipA* | *yehY* | *yhcH* | *ymdB* |
| D42-98-11-12 | CP024881.1 | 98 | 11 | 12 | 1 | 73 | 9 | 9 | 18 |
| D44-98-11-12 | CP026056.1 | 98 | 11 | 12 | 1 | 1 | 9 | 9 | 18 |
| D176-98-11-12 | CP056852.1 | 98 | 11 | 12 | 1 | 1 | 9 | 9 | 18 |
| D265-98-11-12 | CP092463.1 | 98 | 11 | 12 | 1 | 1 | 9 | 9 | 18 |
| D363-98-11-12 | CP135465.1 | 98 | 11 | 12 | 1 | 1 | 9 | 9 | 18 |
| D41-98-11-12 | CP024683.1 | 98 | 11 | 12 | 78 | 1 | 9 | 9 | 18 |
| D156-481-11-4 | CP056527.1 | 481 | 11 | 4 | 56 | 1 | 65 | 5 | 63 |
| D153-519-11-4 | CP056505.1 | 519 | 11 | 4 | 56 | 1 | 65 | 5 | 63 |

**Supplementary Figure S12**

**A**

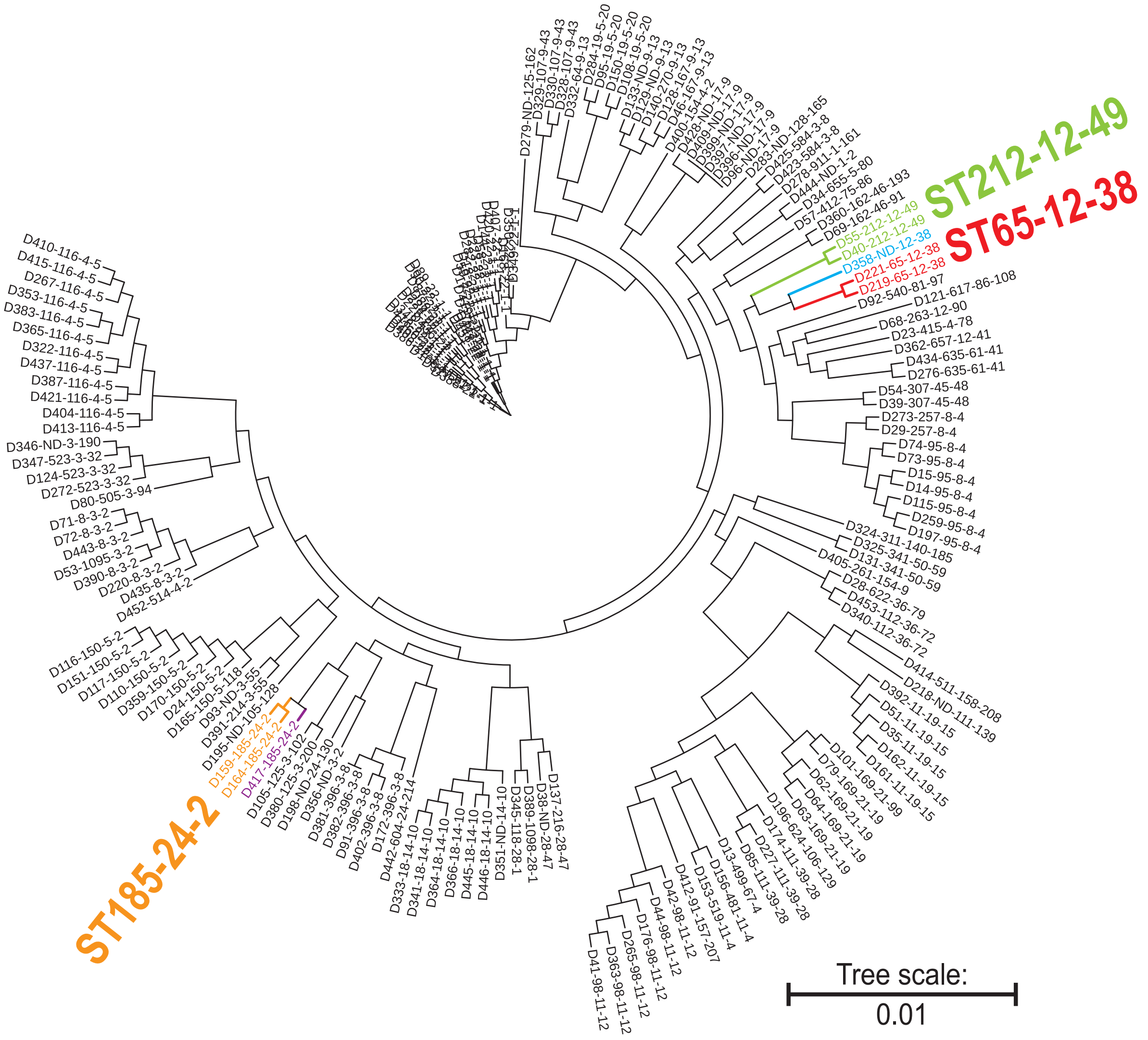


**B**

| | | | Genotypes of the top seven HND genes. | | | | | | |
|---|---|---|---|---|---|---|---|---|---|
| Coding | Accession # | ST | groups_3152 | *nanK* | *iprA* | *mipA* | *yehY* | *yhcH* | *ymdB* |
| D219-65-12-38 | CP070544.1 | 65 | 12 | 38 | 60 | 27 | 72 | 2 | 68 |
| D221-65-12-38 | CP070559.1 | 65 | 12 | 38 | 60 | 27 | 72 | 2 | 68 |
| D358-ND-12-38 | CP132324.1 | ND | 12 | 38 | 1 | 138 | 191 | 2 | 148 |
| D40-212-12-49 | CP024680.1 | 212 | 12 | 49 | 77 | 42 | 51 | 24 | 57 |
| D55-212-12-49 | CP033744.1 | 212 | 12 | 49 | 84 | 42 | 51 | 24 | 57 |
| D68-263-12-90 | CP040698.1 | 263 | 12 | 90 | 6 | 27 | 99 | 25 | 20 |
| D362-657-12-41 | CP135450.1 | 657 | 12 | 41 | 1 | 139 | 193 | 25 | 150 |
| D159-185-24-2 | CP056573.1 | 185 | 24 | 2 | 1 | 6 | 66 | 1 | 7 |
| D164-185-24-2 | CP056622.1 | 185 | 24 | 2 | 1 | 6 | 66 | 1 | 7 |
| D417-185-24-2 | CP162039.1 | 185 | 24 | 2 | 1 | 6 | 210 | 1 | 7 |
| D442-604-24-214 | OW848788.1 | 604 | 24 | 214 | 6 | 152 | 3 | 148 | 7 |
| D198-ND-24-130 | CP059849.1 | ND | 24 | 130 | 1 | 20 | 132 | 2 | 111 |

#### Supplementary Figure S13

A B

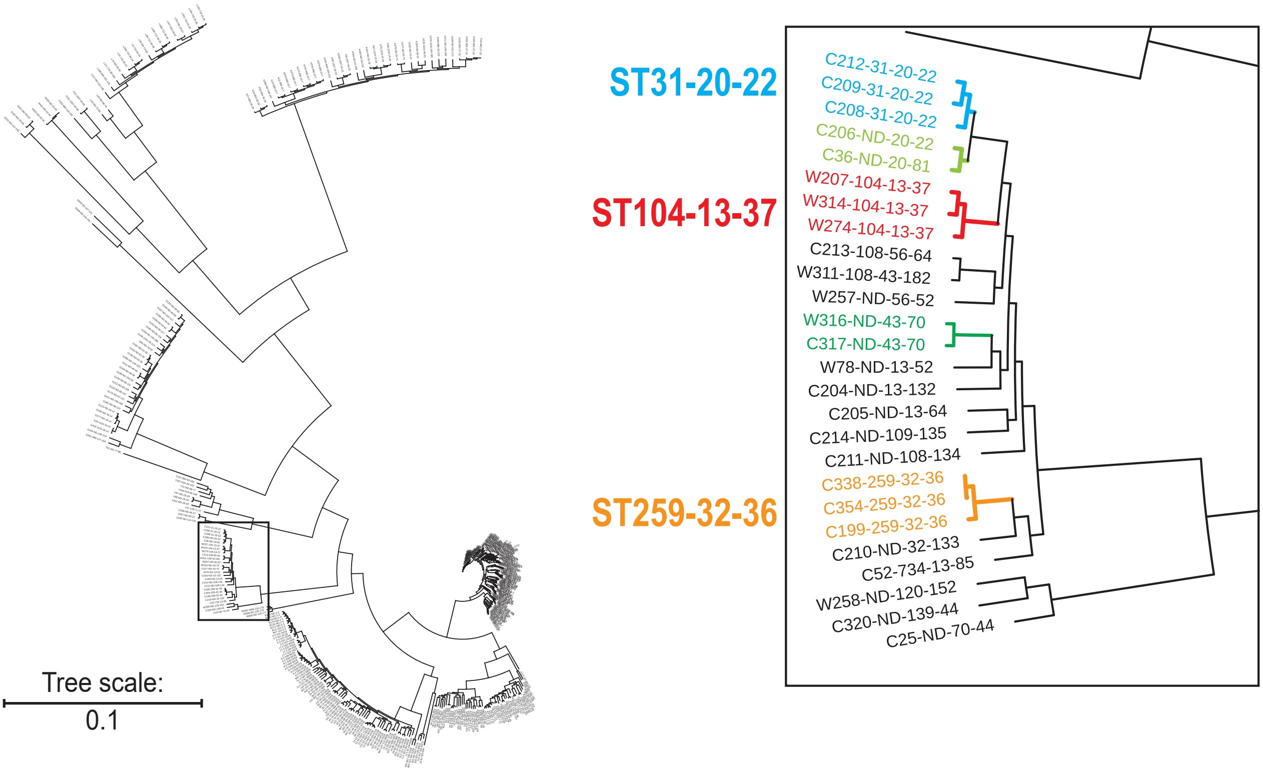


C

| | | | Genotypes of the top seven HND genes. | | | | | | |
|---|---|---|---|---|---|---|---|---|---|
| Coding | Accession # | ST | groups_3152 | *nanK* | *iprA* | *mipA* | *yehY* | *yhcH* | *ymdB* |
| W207-104-13-37 | CP069770.1 | 104 | 13 | 37 | 21 | 34 | 39 | 13 | 114 |
| W274-104-13-37 | CP097324.1 | 104 | 13 | 37 | 21 | 34 | 39 | 13 | 13 |
| W314-104-13-37 | CP101080.1 | 104 | 13 | 37 | 21 | 34 | 39 | 13 | 13 |
| C52-734-13-85 | CP029727.1 | 734 | 13 | 85 | 83 | 77 | 94 | 77 | 88 |
| W78-ND-13-52 | CP044101.1 | ND | 13 | 52 | 89 | 82 | 102 | 13 | 13 |
| C205-ND-13-64 | CP069764.1 | ND | 13 | 64 | 116 | 53 | 134 | 13 | 113 |
| C204-ND-13-132 | CP069763.1 | ND | 13 | 132 | 115 | 26 | 70 | 13 | 13 |
| C208-31-20-22 | CP069779.1 | 31 | 20 | 22 | 15 | 19 | 14 | 17 | 17 |
| C209-31-20-22 | CP069781.1 | 31 | 20 | 22 | 15 | 19 | 14 | 17 | 17 |
| C212-31-20-22 | CP069784.1 | 31 | 20 | 22 | 15 | 19 | 14 | 17 | 17 |
| C206-ND-20-22 | CP069768.1 | ND | 20 | 22 | 15 | 19 | 14 | 17 | 17 |
| C36-ND-20-81 | CP024675.1 | ND | 20 | 81 | 15 | 19 | 14 | 17 | 17 |
| C199-259-32-36 | CP060441.1 | 259 | 32 | 36 | 38 | 26 | 38 | 28 | 33 |
| C338-259-32-36 | CP115032.1 | 259 | 32 | 36 | 38 | 26 | 38 | 28 | 33 |
| C354-259-32-36 | CP126605.1 | 259 | 32 | 36 | 38 | 26 | 38 | 28 | 33 |
| C210-ND-32-133 | CP069782.1 | ND | 32 | 133 | 117 | 98 | 135 | 28 | 33 |
| W311-108-43-182 | CP101066.1 | 108 | 43 | 182 | 59 | 54 | 71 | 29 | 13 |
| W316-ND-43-70 | CP101089.1 | ND | 43 | 70 | 21 | 65 | 80 | 29 | 75 |
| C317-ND-43-70 | CP101092.1 | ND | 43 | 70 | 21 | 65 | 80 | 29 | 75 |

### Supplementary Figure S14

**A**

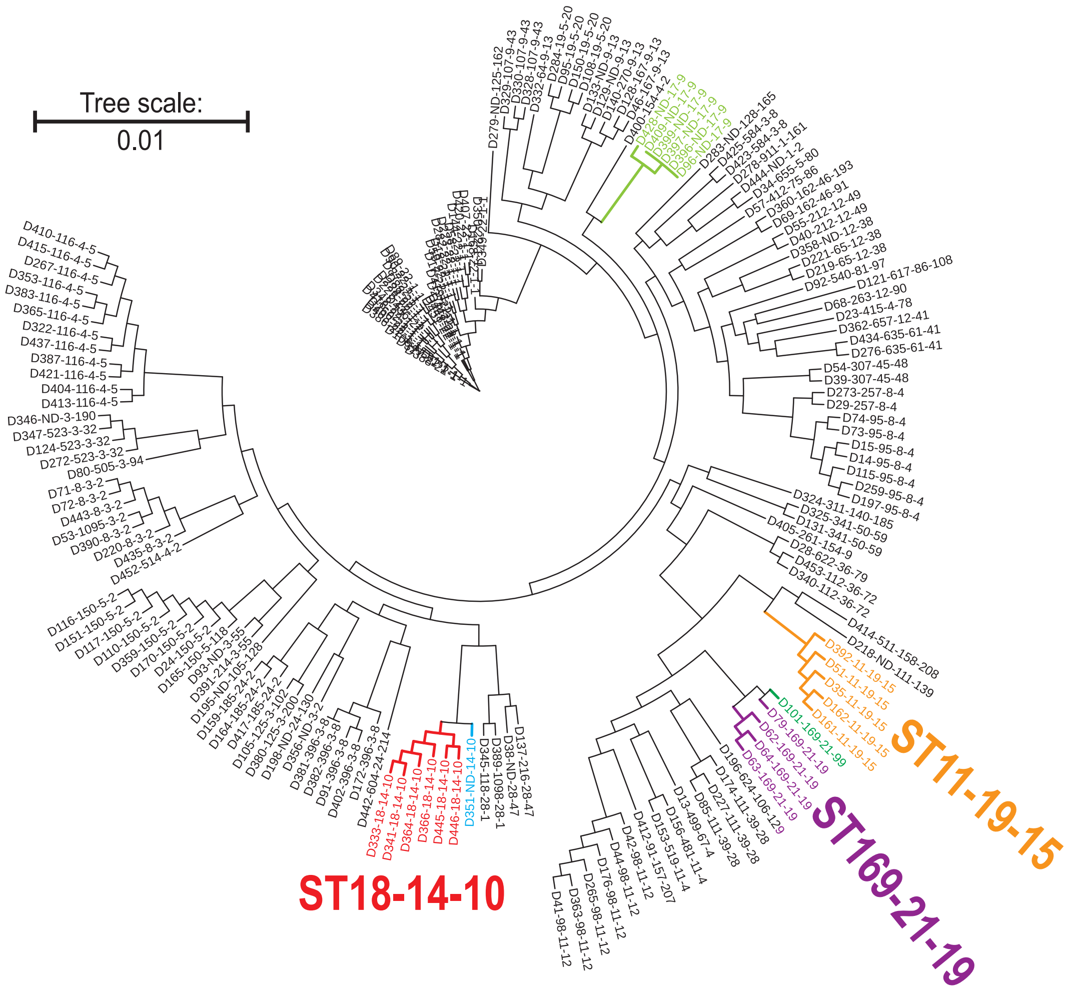


**B**

| Coding | Accession # | ST | Genotypes of the top seven HND genes. | | | | | | |
|---|---|---|---|---|---|---|---|---|---|
| | | | groups_ 3152 | *nanK* | *iprA* | *mipA* | *yehY* | *yhcH* | *ymdB* |
| D333-18-14-10 | CP114564.1 | 18 | 14 | 10 | 6 | 2 | 3 | 14 | 16 |
| D341-18-14-10 | CP115614.1 | 18 | 14 | 10 | 6 | 2 | 3 | 14 | 16 |
| D364-18-14-10 | CP135469.1 | 18 | 14 | 10 | 6 | 2 | 3 | 14 | 16 |
| D366-18-14-10 | CP136047.1 | 18 | 14 | 10 | 6 | 2 | 3 | 14 | 16 |
| D445-18-14-10 | OW849208.1 | 18 | 14 | 10 | 6 | 2 | 3 | 14 | 16 |
| D446-18-14-10 | OW849256.1 | 18 | 14 | 10 | 6 | 2 | 3 | 14 | 16 |
| D351-ND-14-10 | CP125310.1 | ND | 14 | 10 | 6 | 2 | 189 | 14 | 16 |
| D96-ND-17-9 | CP049015.1 | ND | 17 | 9 | 9 | 2 | 12 | 15 | 19 |
| D396-ND-17-9 | CP137703.1 | ND | 17 | 9 | 9 | 2 | 12 | 15 | 19 |
| D397-ND-17-9 | CP137717.1 | ND | 17 | 9 | 9 | 2 | 12 | 15 | 19 |
| D399-ND-17-9 | CP139745.1 | ND | 17 | 9 | 9 | 2 | 12 | 15 | 19 |
| D409-ND-17-9 | CP149135.1 | ND | 17 | 9 | 9 | 2 | 12 | 15 | 19 |
| D428-ND-17-9 | LR134118.1 | ND | 17 | 9 | 166 | 2 | 12 | 15 | 19 |
| D35-11-19-15 | CP024673.1 | 11 | 19 | 15 | 14 | 1 | 13 | 5 | 22 |
| D161-11-19-15 | CP056592.1 | 11 | 19 | 15 | 14 | 1 | 13 | 5 | 22 |
| D162-11-19-15 | CP056595.1 | 11 | 19 | 15 | 14 | 1 | 13 | 5 | 22 |
| D392-11-19-15 | CP137207.1 | 11 | 19 | 15 | 14 | 1 | 13 | 5 | 22 |
| D51-11-19-15 | CP027849.1 | 11 | 19 | 15 | 82 | 1 | 13 | 5 | 22 |
| D62-169-21-19 | CP038653.1 | 169 | 21 | 19 | 27 | 1 | 15 | 1 | 11 |
| D63-169-21-19 | CP038656.1 | 169 | 21 | 19 | 27 | 1 | 15 | 1 | 11 |
| D64-169-21-19 | CP038658.1 | 169 | 21 | 19 | 27 | 1 | 15 | 1 | 11 |
| D79-169-21-19 | CP045555.1 | 169 | 21 | 19 | 90 | 1 | 15 | 1 | 11 |
| D101-169-21-99 | CP052058.1 | 169 | 21 | 99 | 27 | 1 | 15 | 1 | 11 |

Supplementary Figure S15

A

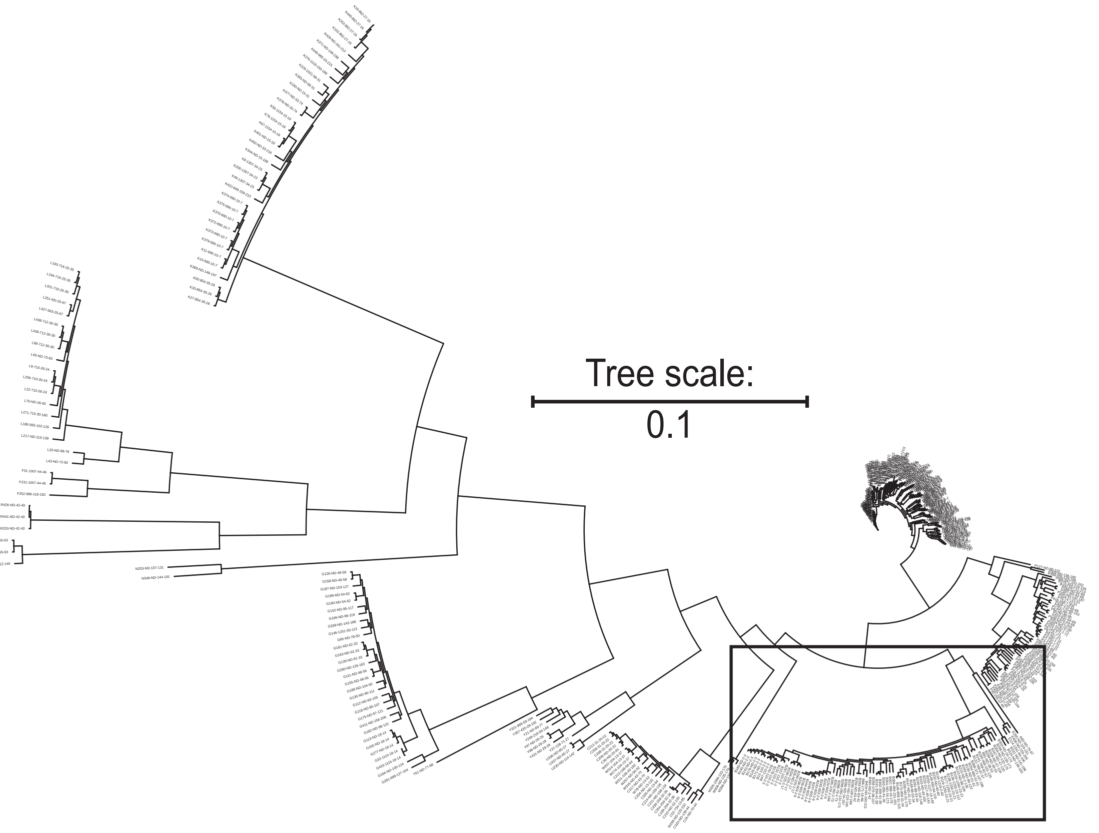

#### Supplementary Figure S15

**B**

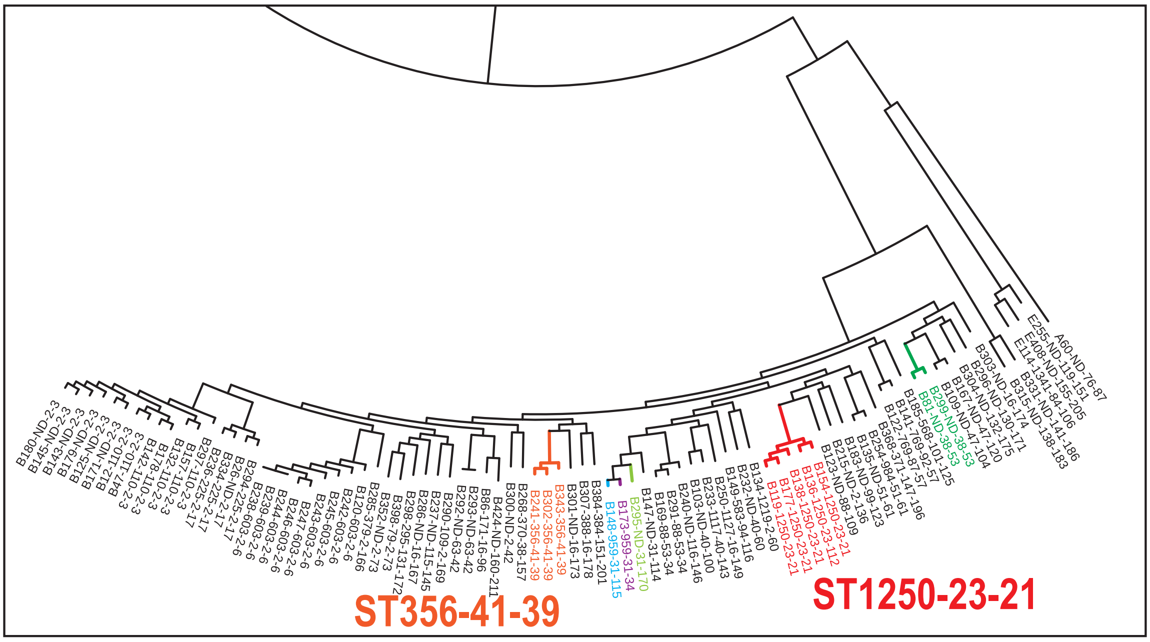


**C**

| | | | Genotypes of the top seven HND genes. | | | | | | |
|---|---|---|---|---|---|---|---|---|---|
| Coding | Accession # | ST | groups_3152 | *nanK* | *iprA* | *mipA* | *yehY* | *yhcH* | *ymdB* |
| B86-171-16-96 | CP046673.1 | 171 | 16 | 96 | 17 | 4 | 103 | 84 | 94 |
| B307-388-16-178 | CP101044.1 | 388 | 16 | 178 | 142 | 62 | 176 | 41 | 14 |
| B250-1127-16-149 | CP080539.1 | 1127 | 16 | 149 | 10 | 112 | 151 | 114 | 124 |
| B286-ND-16-167 | CP099366.1 | ND | 16 | 167 | 7 | 4 | 166 | 41 | 14 |
| B301-ND-16-173 | CP099387.1 | ND | 16 | 173 | 63 | 125 | 171 | 42 | 3 |
| B303-ND-16-174 | CP099389.1 | ND | 16 | 174 | 140 | 126 | 172 | 126 | 14 |
| B119-1250-23-21 | CP055910.1 | 1250 | 23 | 21 | 19 | 21 | 19 | 16 | 25 |
| B138-1250-23-21 | CP056321.1 | 1250 | 23 | 21 | 19 | 21 | 19 | 16 | 25 |
| B154-1250-23-21 | CP056508.1 | 1250 | 23 | 21 | 19 | 21 | 19 | 16 | 25 |
| B177-1250-23-21 | CP056861.1 | 1250 | 23 | 21 | 19 | 21 | 19 | 16 | 25 |
| B136-1250-23-112 | CP056284.1 | 1250 | 23 | 112 | 19 | 21 | 19 | 16 | 25 |
| B148-959-31-115 | CP056409.1 | 959 | 31 | 115 | 10 | 32 | 26 | 27 | 46 |
| B173-959-31-34 | CP056834.1 | 959 | 31 | 34 | 10 | 32 | 26 | 27 | 46 |
| B295-ND-31-170 | CP099379.1 | ND | 31 | 170 | 10 | 32 | 26 | 16 | 14 |
| B147-ND-31-114 | CP056399.1 | ND | 31 | 114 | 10 | 51 | 118 | 57 | 46 |
| B268-370-38-157 | CP092860.1 | 370 | 38 | 157 | 13 | 118 | 158 | 16 | 130 |
| B81-ND-38-53 | CP045771.1 | ND | 38 | 53 | 28 | 43 | 53 | 50 | 40 |
| B299-ND-38-53 | CP099385.1 | ND | 38 | 53 | 138 | 43 | 53 | 50 | 40 |
| B233-1117-40-143 | CP077405.1 | 1117 | 40 | 143 | 49 | 11 | 56 | 63 | 121 |
| B103-ND-40-100 | CP053573.1 | ND | 40 | 100 | 49 | 11 | 56 | 86 | 14 |
| B232-ND-40-60 | CP077300.1 | ND | 40 | 60 | 10 | 107 | 145 | 112 | 120 |
| B241-356-41-39 | CP078598.1 | 356 | 41 | 39 | 30 | 35 | 40 | 39 | 50 |
| B302-356-41-39 | CP099388.1 | 356 | 41 | 39 | 30 | 35 | 40 | 39 | 50 |
| B343-356-41-39 | CP118774.1 | 356 | 41 | 39 | 30 | 35 | 40 | 39 | 50 |

**Supplementary Figure S16**

**A**

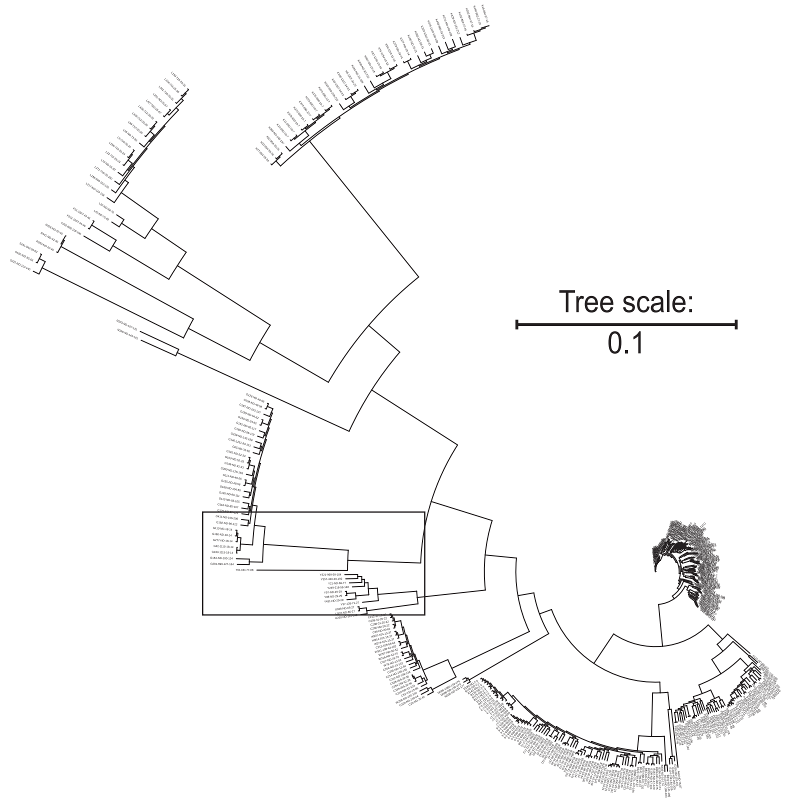


**B**

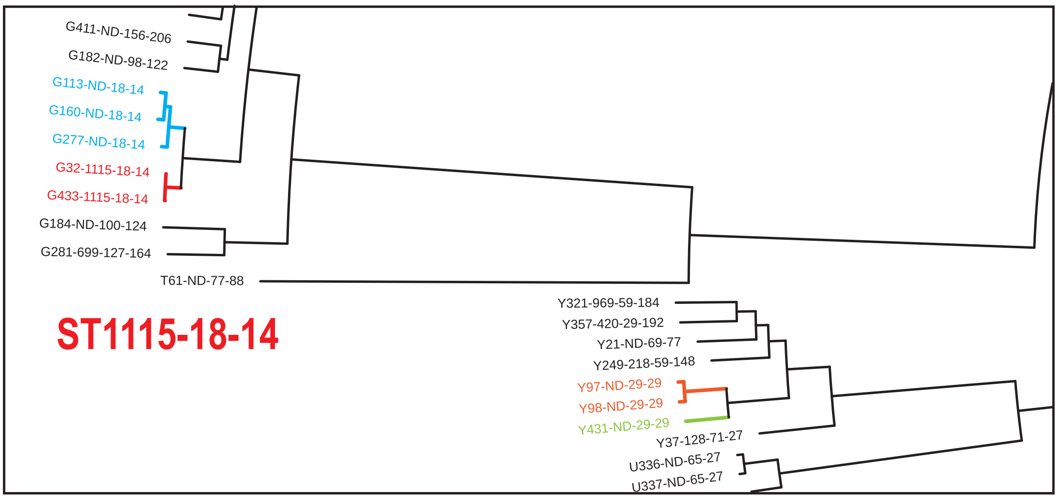


**C**

| | | | Genotypes of the top seven HND genes. | | | | | | |
|---|---|---|---|---|---|---|---|---|---|
| Coding | Accession # | ST | groups_ 3152 | *nanK* | *iprA* | *mipA* | *yehY* | *yhcH* | *ymdB* |
| G32-1115-18-14 | CP023504.1 | 1115 | 18 | 14 | 42 | 41 | 47 | 47 | 21 |
| G433-1115-18-14 | LR699014.1 | 1115 | 18 | 14 | 42 | 41 | 47 | 47 | 21 |
| G113-ND-18-14 | CP055543.1 | ND | 18 | 14 | 51 | 31 | 35 | 34 | 21 |
| G160-ND-18-14 | CP056586.1 | ND | 18 | 14 | 51 | 31 | 35 | 34 | 21 |
| G277-ND-18-14 | CP099084.1 | ND | 18 | 14 | 133 | 31 | 35 | 34 | 21 |
| Y357-420-29-192 | CP132279.1 | 420 | 29 | 192 | 40 | 137 | 190 | 44 | 70 |
| Y97-ND-29-29 | CP049738.1 | ND | 29 | 29 | 33 | 24 | 55 | 52 | 27 |
| Y98-ND-29-29 | CP049739.1 | ND | 29 | 29 | 33 | 24 | 55 | 52 | 27 |
| Y431-ND-29-29 | LR134485.1 | ND | 29 | 29 | 33 | 24 | 214 | 65 | 27 |

#### Supplementary Figure S17

**A**

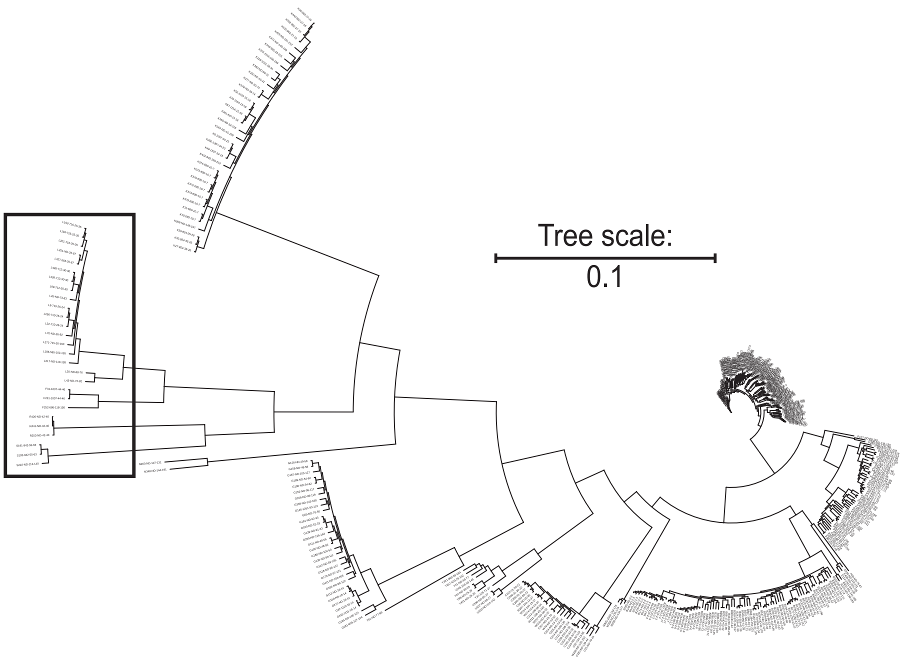


**B**

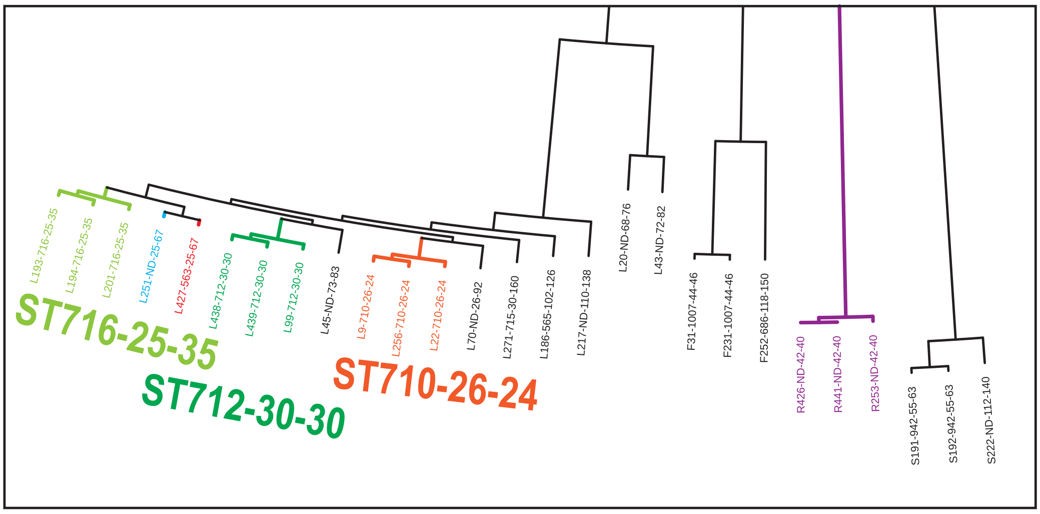


**C**

| | | | Genotypes of the top seven HND genes. | | | | | | |
|---|---|---|---|---|---|---|---|---|---|
| Coding | Accession # | ST | groups_ 3152 | *nanK* | *iprA* | *mipA* | *yehY* | *yhcH* | *ymdB* |
| L427-563-25-67 | JASWFH010000001.1 | 563 | 25 | 67 | 20 | 15 | 73 | 66 | 12 |
| L251-ND-25-67 | CP080958.1 | ND | 25 | 67 | 20 | 15 | 73 | 66 | 12 |
| L193-716-25-35 | CP057630.1 | 716 | 25 | 35 | 20 | 15 | 37 | 38 | 12 |
| L194-716-25-35 | CP057632.1 | 716 | 25 | 35 | 20 | 15 | 37 | 38 | 12 |
| L201-716-25-35 | CP064180.1 | 716 | 25 | 35 | 20 | 15 | 37 | 38 | 12 |
| L22-710-26-24 | CP014070.2 | 710 | 26 | 24 | 32 | 15 | 27 | 31 | 35 |
| L9-710-26-24 | AP024585.1 | 710 | 26 | 24 | 32 | 38 | 27 | 31 | 35 |
| L256-710-26-24 | CP083651.1 | 710 | 26 | 24 | 32 | 38 | 27 | 31 | 35 |
| L70-ND-26-92 | CP041362.1 | ND | 26 | 92 | 46 | 81 | 100 | 82 | 12 |
| L99-712-30-30 | CP050009.1 | 712 | 30 | 30 | 34 | 29 | 33 | 33 | 44 |
| L438-712-30-30 | LT556084.1 | 712 | 30 | 30 | 34 | 29 | 33 | 33 | 44 |
| L439-712-30-30 | LT556085.1 | 712 | 30 | 30 | 34 | 29 | 33 | 33 | 44 |
| L271-715-30-160 | CP096905.1 | 715 | 30 | 160 | 46 | 120 | 160 | 119 | 12 |
| R253-ND-42-40 | CP082833.1 | ND | 42 | 40 | 39 | 36 | 41 | 40 | 51 |
| R426-ND-42-40 | FN543502.1 | ND | 42 | 40 | 39 | 36 | 41 | 40 | 51 |
| R441-ND-42-40 | NC_013716.1 | ND | 42 | 40 | 39 | 36 | 41 | 40 | 51 |

#### Supplementary Figure S18

**A**

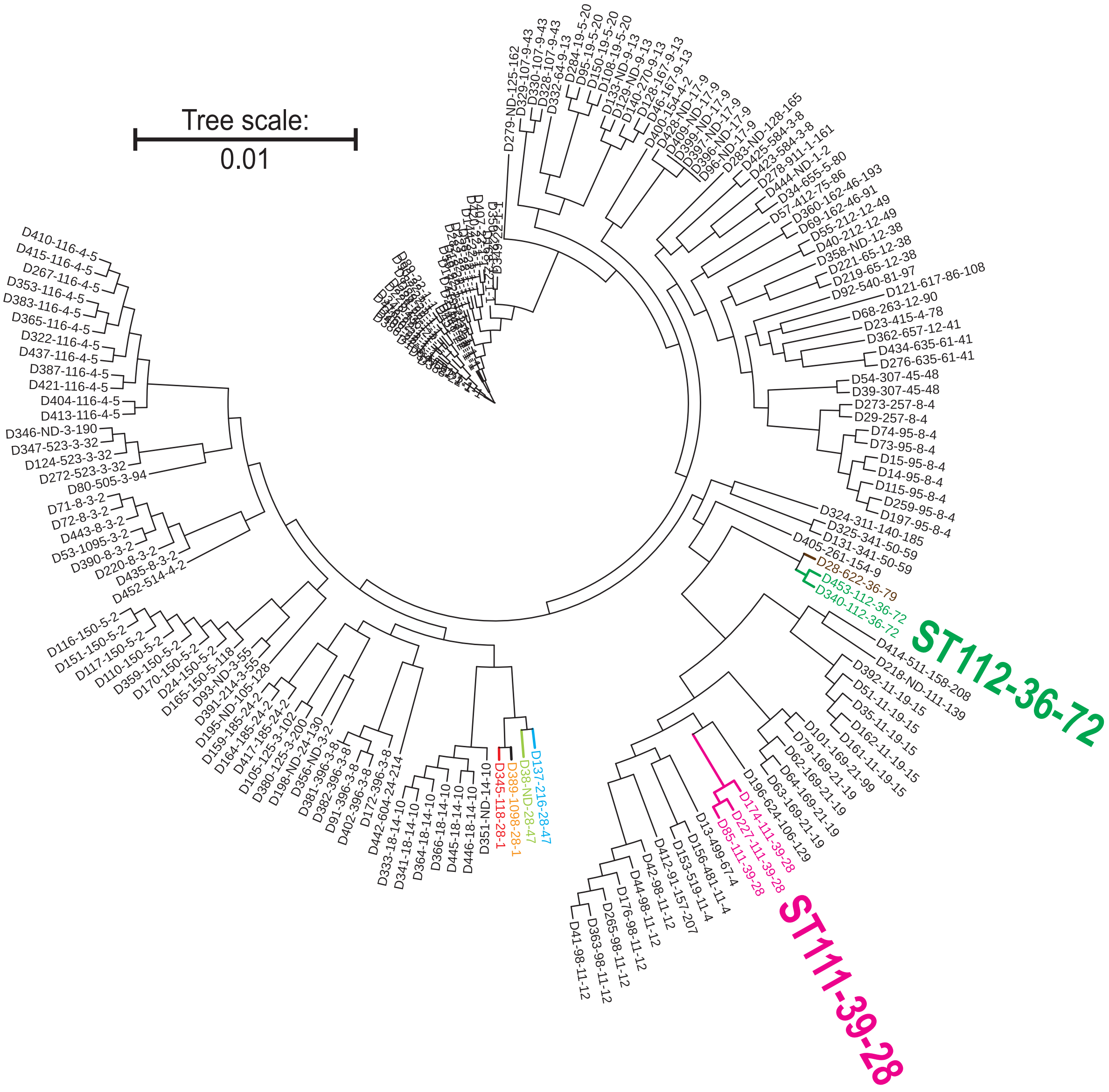


**B**

| Coding | Accession # | ST | Genotypes of the top seven HND genes. | | | | | | |
|---|---|---|---|---|---|---|---|---|---|
| | | | groups_3152 | *nanK* | *iprA* | *mipA* | *yehY* | *yhcH* | *ymdB* |
| D345-118-28-1 | CP119048.1 | 118 | 28 | 1 | 37 | 1 | 3 | 1 | 29 |
| D389-1098-28-1 | CP137195.1 | 1098 | 28 | 1 | 37 | 1 | 3 | 1 | 29 |
| D137-216-28-47 | CP056314.1 | 216 | 28 | 47 | 37 | 1 | 116 | 2 | 29 |
| D38-ND-28-47 | CP024677.1 | ND | 28 | 47 | 76 | 1 | 49 | 2 | 29 |
| D340-112-36-72 | CP115129.1 | 112 | 36 | 72 | 1 | 6 | 44 | 9 | 38 |
| D453-112-36-72 | OW995941.1 | 112 | 36 | 72 | 1 | 6 | 218 | 9 | 38 |
| D28-622-36-79 | CP022151.1 | 622 | 36 | 79 | 1 | 6 | 44 | 9 | 38 |
| D85-111-39-28 | CP046502.1 | 111 | 39 | 28 | 1 | 1 | 32 | 5 | 43 |
| D174-111-39-28 | CP056839.1 | 111 | 39 | 28 | 1 | 1 | 32 | 5 | 43 |
| D227-111-39-28 | CP073043.1 | 111 | 39 | 28 | 1 | 1 | 32 | 5 | 43 |
